# The evolutionary origins of extreme halophilic Archaeal lineages

**DOI:** 10.1101/2019.12.19.883488

**Authors:** Yutian Feng, Uri Neri, Sean Gosselin, Artemis S. Louyakis, R. Thane Papke, Uri Gophna, J. Peter Gogarten

## Abstract

Interest and controversy surrounding the evolutionary origins of extremely halophilic Archaea has increased in recent years, due to the discovery and characterization of the Nanohaloarchaea and the Methanonatronarchaeia. Initial attempts in explaining the evolutionary placement of the two new lineages in relation to the classical Halobacteria (also referred to as Haloarchaea) resulted in hypotheses that imply the new groups share a common ancestor with the Haloarchaea. However, more recent analyses have led to a shift: the Nanohaloarchaea have been largely accepted as being a member of the DPANN superphylum, outside of the euryarchaeota; while the Methanonatronarchaeia have been placed near the base of the Methanotecta (composed of the class II methanogens, the Halobacteriales, and Archaeoglobales). These opposing hypotheses have far-reaching implications on the concepts of convergent evolution (unrelated groups evolve similar strategies for survival), genome reduction, and gene transfer. In this work, we attempt to resolve these conflicts with phylogenetic and phylogenomic data. We provide a robust taxonomic sampling of Archaeal genomes that spans the Asgardarchaea, TACK Group, euryarchaeota, and the DPANN superphylum. In addition, we assembled draft genomes from seven new representatives of the Nanohaloarchaea from distinct geographic locations. Phylogenies derived from these data imply that the highly conserved ATP synthase catalytic/non-catalytic subunits of Nanohaloarchaea share a sisterhood relationship with the Haloarchaea. We also employ a novel gene family distance clustering strategy which shows this sisterhood relationship is not likely the result of a recent gene transfer. In addition, we present and evaluate data that argue for and against the monophyly of the DPANN superphylum, in particular, the inclusion of the Nanohaloarchaea in DPANN.

**Significance Statement:** Many recent analyses have considered large groups of Bacteria and Archaea composed exclusively of environmentally assembled genomes as deep branching taxonomic groups in their respective domains. These groups display characteristics distinct from other members of their domain, which can attract unrelated lineages into those groups. This manuscript evaluates the case of the Nanohaloarchaea, and their inclusion in the DPANN Archaea, through careful analysis of the genes that compose the core of the Nanohaloarchaea. Analyses without inspection of the genes that compose a phylogenomic marker set increases the potential for the inclusion of artifacts and confuses the tree/web of life. Due to horizontal gene transfer and phylogenetic reconstruction artifacts, the placement of divergent archaeal classes into larger groups remains uncertain.

## Introduction

Recent studies discovered several new archaeal lineages in hypersaline environments, including the nanosized Nanohaloarchaea and the methanogenic Methanonatronarchaeia. The exact placement of these lineages within the archaeal phylogeny remains controversial; consequently, the number of independent acquisitions of key adaptations to a halophilic lifestyle remains to be determined. Dissecting the evolutionary relationships between these new lineages and the Haloarchaea may inform on the origins of halophily and the role of genome streamlining. To thrive in extreme hypersaline environments (>150 g/L^-1^), Haloarchaea employ a “salt-in” strategy through the import of potassium ions, in which the intracellular salt concentration equalizes with the external environmental condition (Oren, 2008). This acts to balance the cellular osmotic pressure but also has caused significant changes in amino acid usage, leading to an overabundance of acidic residues, aspartate and glutamate (D/E) in all Haloarchaea (Lanyi, 1974; Madern et al., 2000).

The evolutionary origins of the Nanohaloarchaea have remained uncertain since their discovery (Ghai *et al*., 2011; Narasingarao *et al*., 2012). The composition of their proteome indicates that Nanohaloarchaea also use the “salt-in” strategy similar to Haloarchaea (Narasingarao *et al*., 2012). It was originally suggested that the Nanohaloarchaea are euryarchaeota that form a clade with the Haloarchaea, based on phylogenies of the 16S rRNA gene and ribosomal proteins (Narasingarao *et al*., 2012; Petitjean *et al*., 2014). Additional data obtained from individual cells via cell sorting followed by genome amplification and 16S rRNA sequencing analysis confirmed the original observations of the Nanohaloarchaea as a sister taxon to the Haloarchaea (Zhaxybayeva *et al*., 2013). More recently, based on analyses of concatenated conserved protein sequences, the Nanohaloarchaea were placed in a group together with similarly nanosized organisms, the Diapherotrites, Parvarachaeota, Aenigmarchaeota, and Nanoarchaeota, forming the DPANN superphylum (Andrade *et al*., 2015; Castelle *et al*., 2015; Rinke *et al*., 2013).

Past analyses of this superphylum (Brochier-Armanet *et al*., 2011; Petitjean *et al*., 2014; Raymann *et al*., 2014; Williams *et al*., 2015) suggested that the DPANN grouping may not reflect shared ancestry but rather an artifact due to long branches and/or small genomes. However, more recent analyses supported a monophyletic DPANN clade (Williams et al., 2017). Aouad *et al*. performed a multi-locus analysis using various models, which did not include DPANN sequences, and placed the Nanohaloarchaea with the Methanocellales and the Haloarchaea with the Methanomicrobiales (Aouad *et al*., 2018); *i.e.*, the Nanohaloarchaea were recovered as a member of the euryarchaeota, but not as a sister-group to the Haloarchaea. We note that a similar controversy surrounds the phylogenetic position of the Nanoarchaeota.

*Nanoarchaeum equitans* was first considered a representative of a new deep branching archaeal phylum (Huber *et al*., 2002), *i.e.,* an archaeon not a member of the euryarchaeotes or crenarchaeotes. However, later analyses of ribosomal proteins, phylogenetically informative HGTs, and signature genes led to the conclusion that *N. equitans* may represent a fast-evolving euryarchaeote instead of an early branching novel phylum (Brochier *et al*., 2005; Dutilh *et al*., 2008; Urbonavičius *et al*., 2008). Several more recent analyses placed the Nanoarchaeota inside of the DPANN (*(Adam et al., 2017; Dombrowski et al*., 2019; Spang *et al*., 2017), reflecting the ongoing controversy in the phylogenetic placement of these groups.

Recently, another group of extreme halophiles, the Methanonatronarchaeia (also spelled as Methanonatronarcheia), were discovered and predicted to also use the “salt-in” strategy (Sorokin *et al*., 2017). Initial multi-locus phylogenetic analyses placed these methanogenic halophiles in a monophyletic clade with the Haloarchaea, suggesting they are an evolutionary intermediate between methanogens and modern halophiles. However, several recent studies have contested this placement: a multi-locus dataset placed the Methanonatronarchaeia basal to a superclass named Methanotecta, a group that includes the Archaeoglobales, class II methanogens and Haloarchaea (Adam *et al*., 2017; Aouad *et al*., 2019; Martijn *et al*., 2020).In addition, to the three extreme halophiles mentioned, the recently characterized Hikarchaeia has been identified as a non-halophilic sister-group to the Haloarchea (Martijn *et al*., 2020). Temporal analysis of the Hikarchaeia divergence from the Haloarchaea may shed light on the genomic events that prelude the Haloarchaea’s adaptation to hypersalinity (see discussion).

Several conclusions can be drawn from these latter results with regard to adaptation to a halophilic lifestyle, most note-worthy of which is the convergent evolution of the “salt-in” strategy among these three lineages. Independent adaptation to hypersalinity in extreme halophiles is certainly a viable evolutionary hypothesis; this is seen in the case of the *Salinibacter* and *Salinicoccus* (Mongodin *et al*., 2005). However, if Nanohaloarchaea, Haloarchaea, and Methanonatronarcheia form a monophyletic group, as seen with some analyses of 16S rRNA and ribosomal proteins, the hypothesis of common ancestral origins can more easily account for the evolutionary development of the salt-in strategy.

The evolutionary relationships of the three extreme halophilic archaeal lineages remain unresolved; Figure 1 summarizes the current controversies. This lack of resolution can, at least in part, be due to biases that are known to complicate phylogenetics. The genomes of the Methanonatronarchaeia and Nanohaloarchaea are comparatively small with average genome sizes of <2.1Mb and ∼1.1 Mb, respectively. Furthermore, most genome entries in public databases are incomplete. The Haloarchaea are known to be highly recombinogenic (Boucher *et al*., 2004; Méheust *et al*., 2018; Mohan *et al*., 2014; Naor *et al*., 2012; D. Williams *et al*.,2012) and are physically associated with at least some of the Nanohaloarchaea (Andrade *et al*., 2015; Cono *et al*., 2019; Hamm *et al*., 2019).

**Fig 1.**
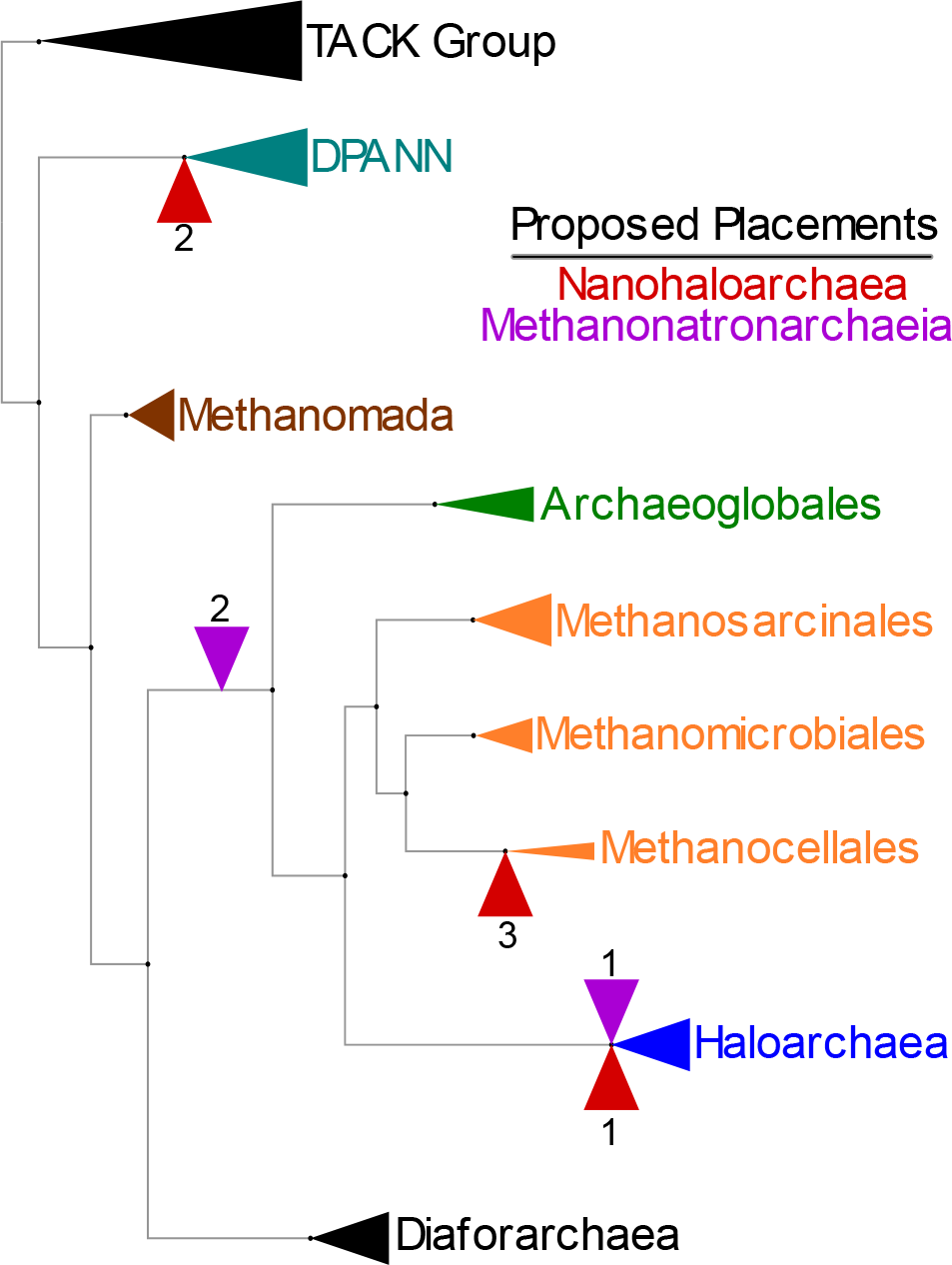
Summary of proposed placements of halophilic lineages mapped on an Archaeal reference tree. This reference tree mostly depicts the positions of various euryarchaea. Individual taxa have been collapsed into higher taxonomic groups. The red (R) indicators represent the different placements proposed for the Nanohaloarchaea, while the purple (P) indicators are used for the Methanonatronarchaeia. Sources for each placement: R1 (Narasingarao *et al*., 2012), R2 (Andrade *et al*., 2015), R3 (Aouad *et al*., 2018); P1 (Sorokin *et al*., 2017), and P2 (Aouad *et al.,* 2019).

Phylogenies based on the concatenation of many genes face many problems: 1) Genes have different evolutionary histories (e.g. duplication and transfer) and forcing the histories of all the genes on a single tree does not reflect the complex evolutionary history of the genomes (Lapierre *et al*., 2014). In particular, genes acquired from outside the group under consideration may create a strong signal for placing the recipient of the transferred gene at the base of the group. 2) Genes experience differing levels of purifying selection, especially between different lineages. This can lead to long branch attraction (LBA) artifacts (Felsenstein, 1978), even if the individual genes evolved along the same history as the host species (Philippe *et al*., 2005). 3) Substitution bias may create convergent signals in unrelated groups.

The work reported here was guided by the hypothesis that the phylogenetic reconstruction of a single, slowly evolving gene might be more robust against artifacts of phylogenetic reconstructions compared to analyses that are based on large sets of genes that may represent different evolutionary histories, include missing data, and contain genes with high substitution rates. We reconstruct single gene alongside multi-locus phylogenies to correct for these sources of bias and to critically assess the evolutionary relationships of the Haloarchaea, Nanohaloarchaea, and Methanonatronarchaeia. We also cluster and dissect the evolutionary relationships of the gene families in the Nanohaloarchaeal core genome, using a gene family clustering technique.

The ATP synthase catalytic and non-catalytic subunits, AtpA and AtpB, represent extremely slow evolving genes (Gogarten, 1994) conserved throughout Archaea and are among the slowest evolving genes in cellular organisms (Table S5). The evolution of these subunits may be slow enough to ameliorate rate signal bias and minimize compositional heterogeneity that otherwise plague reconstructions that includes DPANN and Haloarchaeal sequences. ATP synthase subunits have been used successfully as a phylogenetic marker for large scale reconstructions (Gogarten & Taiz, 1992); however, a drawback of the ATPases is that they are known to have been transferred between divergent phyla (Olendzenski *et al*., 2000). Recently, Wang *et al*. convincingly showed the transfer of this operon lead to the adaptation of Thaumarchaeota to more acidic environments (Wang *et al.,* 2019). The same authors drew a similar conclusion when the Nanohaloarchaea-Haloarchaea sister-group was recovered, which the authors interpreted as suggesting HGT of the ATPase genes in the Nanohaloarchaea-Haloarchaea. To shed light on this HGT hypothesis, we cluster and correlate the gene families in the Nanohaloarchaea and contrast the position of the ATPase genes in these clusters to the same genes in the Thaumarchaeota. We also provide a more robust sampling of the Nanohaloarchaea; we include seven newly sequenced and assembled nanohaloarchaeal genomes together with existing genomes mined from the NCBI database. Robust sampling of the taxa of interest, like the one offered here, has the potential to improve the recovery of evolutionary relationships without adding more sites (genes) (Graybeal, 1998).

In maximum likelihood and Bayesian phylogenies, we find that the Nanohaloarchaea group robustly with the Haloarchaea in the single gene phylogenies, while the Methanonatronarchaeia were placed as a deeper branching euryarchaeal lineage, most likely at the base of the Methanotecta superclass. In large, concatenated datasets, we recover a monophyletic DPANN (including the Nanohaloarchaea). We also provide evidence that the ATPase genes have likely not been transferred in the case of the Nanohaloarchaea-Haloarchaea, and contrast this specific relationship with the clearly transferred ATPases in the Thaumarchaeota.

## Results

### Increased genomic representation of the Nanohaloarchaea

We obtained five new Nanohaloarchaea single amplified genomes (SAGs) from solar salterns in Spain and two metagenome assembled genomes (MAGs) from Israel. The summary statistics and accompanying information of these genomes can be found in Table S2. While the SAGs are of poor assembly quality and completeness, enough genes were recovered from them for phylogenomic analyses; furthermore, they unequivocally group with the other Nanohaloarchaea in all analyses. These seven genomes expand the number of Nanohaloarchaea assemblies available for analyses (18 at time of writing). Total average nucleotide identity (tANI) was used to delineate taxonomy amongst the newly described Nanohaloarchaea. Figure S2 is a distance-based tree calculated from corrected tANI distance (see Figure S1 for distance matrix) between the previously described and newly described Nanohaloarchaea. Using conservative cutoffs, it appears that SAGs SCGC AAA188-M06 and M04 may belong to the genus *Ca.* Nanosalina. SAG M21 seems to be a member of *Ca*. Nanosalinarium, while the remaining new genomes (SAGs and MAGs) do not belong to any previously described candidate genera.

The genome described as Nanohaloarchaea archaeon PL-Br10-U2g5 (Vavourakis *et al*., 2016) was likely miss-identified as a Nanohaloarchaeon. We find that this strain unequivocally groups within *Halorubrum* species in ribosomal (protein and rRNA), whole genome, and single gene phylogenies (Figure S2).

While the fragmented nature of the SAGs is useful for phylogenetic analyses, there are not enough genes to paint a comprehensive picture of their inferred metabolisms; the focus of the metabolic analyses was therefore centered on the two MAGs described above as they are almost complete. The two MAGs, M322 and AT22, have metabolic capabilities comparable to previously described Nanohaloarchaea (Narasingarao *et al*., 2012). Both MAGs are deficient for enzymes in their nitrogen incorporation and lipid biosynthesis pathways. Both genomes encode key enzymes involved in glycolysis and sugar metabolism; the presence of a number of sugar dehydrogenases indicates a possible fermentative lifestyle. AT22 encodes a membrane bound domain and the jellyroll fold LamG, which have been implicated in host cell interactions in DPANN archaea (Golyshina *et al*., 2017; Hamm *et al*., 2019). The comprehensive list of genes in their respective genomes (including the SAGs) described here are provided in Table S3.

### Phylogenetic placement of halophilic lineages

To shed light on the evolutionary origins of the Methanonatronarchaeia and the Nanohaloarchaea, we have produced three sets of trees from distinct markers that contain differing phylogenetic signals. A marker set composed of the AtpAB proteins (Figure 2, S3) was used to calculate phylogenies; these are slowly evolving single copy genes. The phylogenies calculated from this marker set were compared to phylogenies calculated by large concatenates: a concatenate of 44 ribosomal proteins (Figure S6), and a concatenate of 282 genes calculated to be within the core genome of the Nanohaloarchaea (Figures 5, S7,9). All three tree sets contain >150 taxa, representing Archaea that span the euryarchaeota, TACK group (including Asgard archaea), and the candidate DPANN superphylum. The phylogenies are depicted as rooted with the TACK Group, but should be considered as **unrooted**, as the root of the Archaeal tree remains an open question with the emergence of Eukaryotes from the Archaea likely having rendered them paraphyletic (Fournier & Poole, 2018; Gribaldo *et al*., 2010; Spang et al., 2018; Williams *et al*., 2020; Williams *et al.,* 2013). In addition, a recent study places the root inside of the euryarchaeota (Raymann *et al*., 2015). However, the placement of the Archaeal root does not impact the conclusions drawn from our phylogenies, presuming the Archaeal root is placed outside of the euryarchaeal crown group.

**Fig 2.**
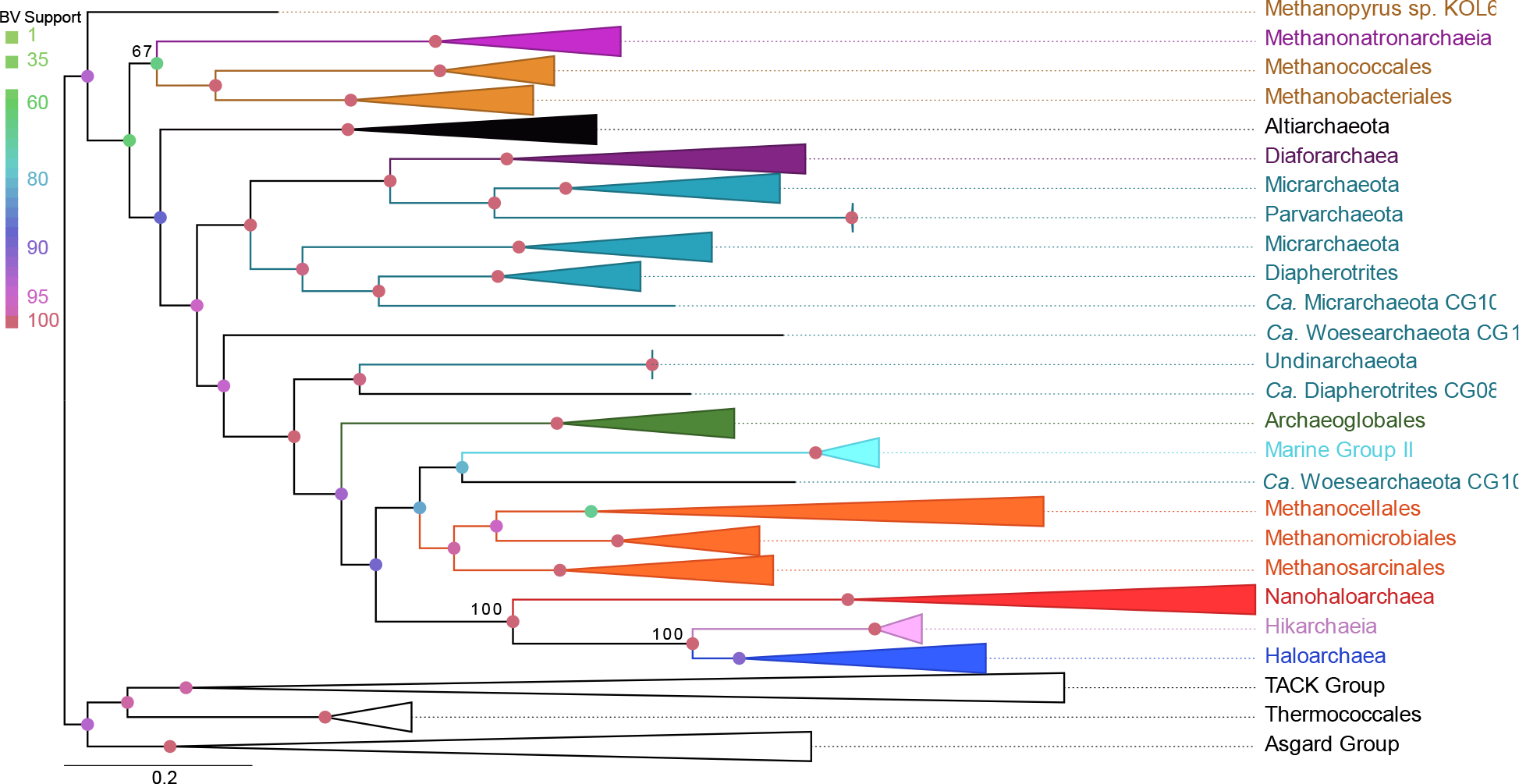
Maximum likelihood phylogeny calculated from AtpAB proteins. The depicted tree contains most features of the other calculated ATP synthase phylogenies. Several taxa were collapsed into higher taxonomic ranks. Important taxa including the halophilic lineages and DPANN (teal) sequences have been colored; *Nanohaloarchaea* (red), *Haloarchaea* (blue), *Methanonatronarchaeia* (purple), Methanomada (brown), Methanotecta methanogens (orange), and the Hikarchaeia (magenta). The tree is drawn as rooted by the TACK Group but should be considered as unrooted. This tree was calculated using the LG+C60 model.

### ATP synthase catalytic and non-catalytic subunit phylogenies

The ATP synthase catalytic and non-catalytic subunits are slow evolving, essential genes. Single protein phylogenies of these subunits may ameliorate LBA and deletion-transfer-loss (DTL: evolutionary conflicts driven by gene transfer, loss, or deletion that may mislead the interpretation of a phylogeny) conflicts; both of which have plagued large scale, multi-locus attempts at reconstructing the Archaeal phylogeny. A drawback of the ATP synthase is that it has been suggested to have been horizontally transferred, and thus its phylogeny, although less prone to artifacts of phylogenetic reconstruction, may not represent the whole genome (see discussion below). Maximum likelihood phylogenies (see Table S4) of the AtpA and AtpB proteins were created using site-homogeneous and site-heterogeneous substitution models. All tree reconstructions based on the original, unaltered multiple sequence alignments confidently placed the Nanohaloarchaea as a sister-group to the Haloarchaea (≥ 91 Bootstrap Value (BV)). A representative example tree constructed with the AtpA+B concatenate is shown below (Figure 2). Nanohaloarchaea and Haloarchaea are grouped together and are positioned inside the Methanotecta. Typically, Haloarchaea are often seen as a sister-group to the Class II methanogens (group including the Methanosarcinales, Methanomicrobiales, and Methanocellales), in ATPases subunit phylogenies this relationship was interrupted by the placement of other archaeal groups: in case of the AtpA+B concatenate and AtpA, the Nanohaloarchaea-Haloarchaea sistergroup is separated from the Class II methanogens by Marine Group II archaea and a Woesearchaeon (Figure 2, Table S4, Figure S3a). In case of the AtpB protein the Nanohaloarchaea-Haloarchaea sistergroup is also sometimes (dependent on type of substitution model, % of sites retained, and alterations to the alignment matrix, i.e., recoding) recovered at the base of the Methanotecta (a group including the class II methanogens and the Archaeoglobales) (Table S4, Figure S3b).

Curiously, the Methanonatronarchaeia are placed as a deeper branching euryarchaeal lineage, suggesting either gene transfer or convergent evolution in regard to the extreme halophilic “salt-in” strategy. In only one analysis did all three lineages group together, with poor support inside the Methanotecta, (Table S4, LG+C50 AtpA d4). The newly described Hikarchaeia’s (non-halophilic sister-group of the Haloarchaea; Martijn *et al*., 2020) sisterhood with the Haloarchaea is recovered in the AtpA and AtpA+B sequences (Figure 2, Figure S3a). In the case of the AtpB tree, the Hikarchaeia emerged from within the Haloarchaea (Figure S3b), but with marginal support (BV= 68). In all of these ATPase subunit phylogenies the Nanohaloarchaeota branch before the split between Haloarchaea and Hikarchaeia.

The placement of the remaining DPANN taxa (i.e., without Nanohaloarchaea) appears erratic. However, it is worth noting the groups considered as members of DPANN fail to form a monophyletic clade in all of the ATPase based trees and the branches breaking the DPANN group apart are supported by high BVs (Table S4).

### Compositional bias

Compositional bias in encoded amino acids can generate artifacts in large, domain-wide phylogenies (Aouad *et al.,* 2018; Aouad *et al.,* 2019). However, due to the slow rate of evolution in these ATPase subunits, compositional bias has been minimized. A chi-squared test of composition for both protein alignments revealed only 10% and 6% of taxa fail the composition test in AtpA and AtpB sequences, respectively. None of the sequences that failed this composition test belong to a member of the halophilic lineages barring one sequence that belonged to a Nanohaloarchaeon with an incompletely sequenced *atpA.* To minimize compositional bias, both alignments were recoded into 4 and 6 Dayhoff groups (Susko & Roger, 2007). These recoded alignments were used to create maximum likelihood and Bayesian phylogenies, which mostly recapitulated the groupings discussed earlier (Table S4). The only difference was that in several instances Methanonatronarchaeia moved either to the base of the Methanotecta, grouped with the Haloarchaea and Nanohaloarchaea, or with the TACK group.

A reason for bias in extreme halophilic lineages is an acidic proteome, *i.e.,* increased presence of aspartic and glutamic acid (D/E) in their protein sequence. This may lead to “compositional attraction”, where those taxa that have an abundance of D/E sites are more likely to cluster together in a phylogeny. Sites that contained a conserved D/E residue among the Haloarchaea, Nanohaloarchaea, and the Methanonatronarchaeia were deleted from the AtpA and AtpB alignments. Maximum likelihood phylogenies were created from these new alignments (Table S4), and the topology discussed above was recovered, albeit with lower support due to the loss of phylogenetically informative sites. We recognize that this method limits compositional attraction, but cannot completely rule out the possibility other residues have evolved independently in similar hypersaline environments.

### Suitability of the ATP synthase subunits as a phylogenetic marker

Although the ATP synthase subunits are an attractive phylogenetic marker, these genes have been shown to be horizontally transferred (Wang *et al*., 2019), sometimes between distantly related organisms (Lapierre, *et al*., 2006; Olendzenski *et al*., 2000), and are presumably adaptive. Wang *et al*. has recently shown, convincingly, that Thaumarchaeota have adapted to more acidic environments, with a gene transfer of V-Type ATPases from other acidophilic archaea. A similar transfer has also been suggested for the Nanohaloarchaea, which would complicate the interpretation of the ATPase phylogenies. To shed light on whether this is the case, we regressed and correlated the pairwise distance matrices of the gene families in the Nanohaloarchaea and the Thaumarchaeota (using a modification of the approach described in (Rangel *et al*., 2019; Rangel *et al*., 2020) see methods for details). Two analyses were performed with this gene family distance method, with different marker sets. The GTDB Archaeal 122 marker set (Parks *et al*., 2018) with the addition of ATPase genes was used to compare the correlation of gene families within the Thaumarchaeota and the Nanohaloarchaea. These markers have been found to produce consistent phylogenies, and should be commonly found in most Archaeal genomes, thus they are appropriate for making comparisons between two groups. Relationships (via correlation distance 1-r^2^) between gene families in the Nanohaloarchaea and Thaumarcheota were calculated, and further ordinated using nMDS (non-metric multi-dimensional scaling) and statistically evaluated using a categorical Mantel test at the 95% confidence level (Figure 3).

**Fig 3.**
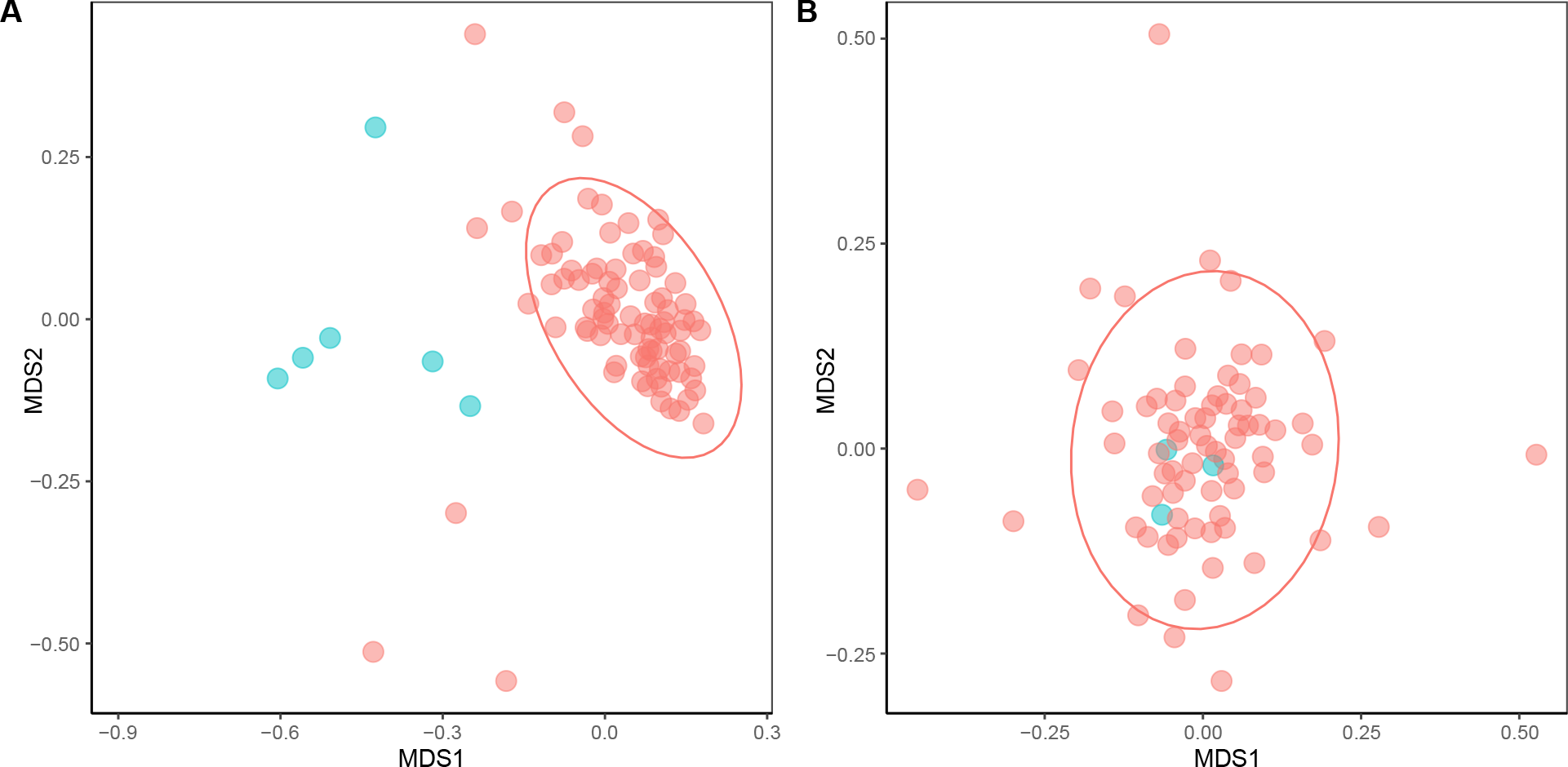
nMDS plots of the gene families in the Thaumarchaeota and Nanohaloarchaea. Shows the ordination of various gene families (from the Archaea 122 marker set) in the Thaumarchaeota and Nanohaloarchaea. A categorical Mantel test with two defined categories, ATPase genes (colored in blue) and non-ATPase genes (colored in red), was used to determine significance with the 95% confidence ellipse. A) The gene families in the Thaumarchaeota, the ATPase genes clearly fall outside of the 95% confidence ellipse, with a p = 0.001. B) The gene families in the Nanohaloarchaea, where the ATPase genes clearly fall inside the ellipse, with a p = 0.182.

The ordination plots (Figure 3) clearly show the contrast of the relative evolutionary trajectories of the ATPase genes in the Thaumarchaeota versus Nanohaloarchaea. In the case of the Thaumarchaeota, whose ATPase genes are known to be transferred (Wang *et al.,* 2019), the ATPase genes clearly fall outside of the 95% confidence ellipse. The 95% confidence ellipse, in this analysis, comprises most of the gene families belonging to a common evolutionary trajectory and are likely not the result of a recent horizontal gene transfer between divergent species. The stark contrast between the ATPase genes in these two groups, highlights the difference in circumstance around the evolutionary trajectory of the ATPase genes relative to other gene families in both groups. For reference, this analysis was also performed on subsets of the Thaumarchaeota genomes (Wang *et al*., 2019); one subset containing the acidophiles (received V Type ATPases via HGT) and another subset containing only the neutrophilic Thaumarchaeota (vertically inherited A type ATPases). The acidophilic Thaumarchaeota’s ATPases also stands out as atypical in this analysis (transfer detected, p-value = 0.024; Figure S14a), while the ATPases in the neutrophiles falls comfortably inside the 95% confidence ellipse (p-value = 0.576).

In addition to the ordinations, the correlations between the gene family distance matrices and their associated clustering diagrams for both the Archaeal 122 and the 282 core gene marker sets revealed, that in the Nanohaloarchaea (Figures. 4, S4), the ATPase genes fall into a large cluster of genes (Figure 4, genes highlighted by purple rectangle from 282 core genes of the Nanohaloarchaea). Every gene family enclosed by these purple rectangles, including the AtpAB genes, share a broadly similar evolutionary trajectory. This cluster is also clearly distant to other large clusters (i.e., genes enclosed by the blue rectangle in Figure 4) and the genes located on deep, long branches that fail to form a significantly large cluster. The genes on deep, long branches in these clustering diagrams represent those that likely have been horizontally transferred or follow an unconventional (i.e., not strictly vertical) evolutionary history, reflected in the pairwise distance matrices of the gene family. The genes within the large clusters share a similar evolutionary history, which might be explained with predominantly vertical inheritance. In the gene families of the Thaumarchaeota (Figure S5), the ATPase genes (which were identified as having been transferred, Wang *et al.,* 2019) form a separate cluster distant from three other clusters. The Thaumarchaeota ATPase subunits likely share an evolutionary trajectory with each other that differs from the trajectories in the other two clusters and other individual genes on long branches. A description of this clustering implementation can be found in the methods section, and in an interactive Jupyter notebook script with instructions (https://github.com/Gogarten-Lab-Team/NanoH_GBE_2020).

**Figure 4.**
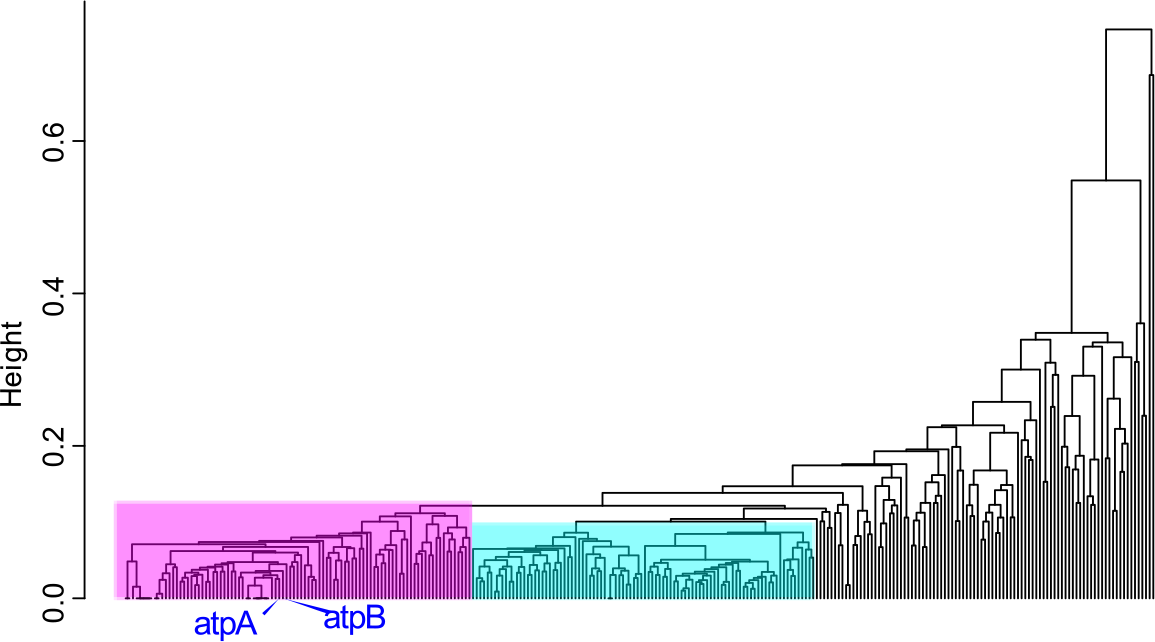
**Clustering diagram of 282 gene families that form the core to the Nanohaloarchaea**, clustered by the pairwise correlation between distance matrices calculated for individual gene families. Families clustered together share similar (although not identical) evolutionary trajectories as assessed by their distance matrices calculated using maximum likelihood models (see materials and methods). Gene families enclosed by the rectangles share broadly similar evolutionary trajectories (with the same members of their cluster), and likely not have been transferred between divergent lineages, while gene families on deep, long branches likely have an unconventional evolutionary trajectory. Subdivisions of the Large Core supermatrix were defined using the clusters (rectangles) in the dendrogram, called the Left (gene families enclosed by the purple rectangle), Right (blue rectangle), and Center (a combination of Left and Right clusters). The blue tip labels indicates where the AtpAB genes fall in the clusters.

**Fig 5.**
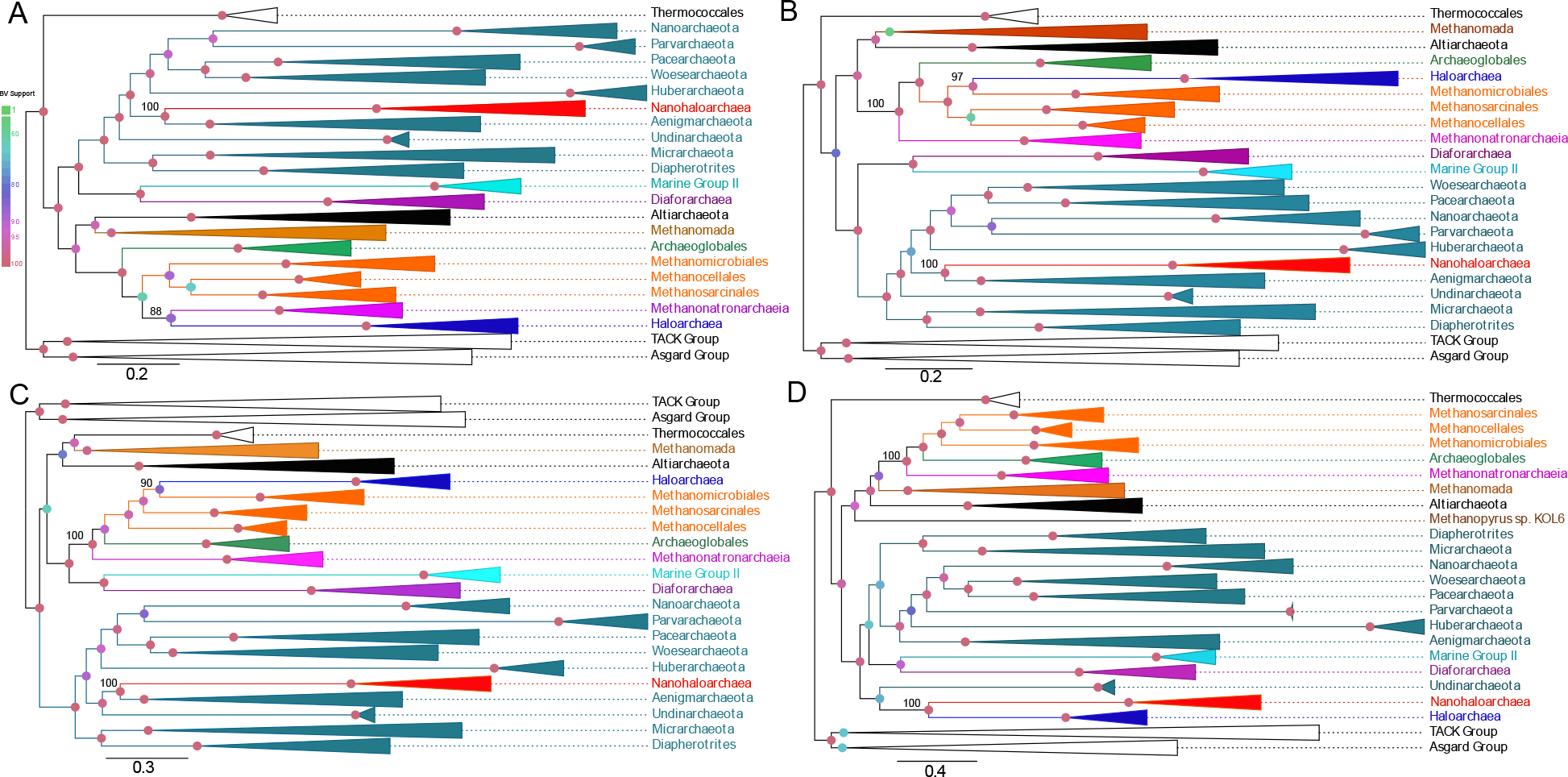
Maximum likelihood phylogenies of Archaeal Large Core genome supermatrices. All phylogenies were calculated with the LG+C60 mixture model A) Maximum likelihood phylogeny calculated using all 282 gene families. B) Phylogeny calculated using the Center supermatrix. C) Phylogeny calculated using the Left supermatrix (95 gene families). D) Phylogeny calculated using the Right supermatrix (94 gene families). Colored node circles indicate bootstrap support value magnitude.

The 282 core gene families of the Nanohaloarchaea were also ranked on their speed of evolution; based on their average slopes of pairwise distance matrices (1 gene family vs. the 281 other gene families) and their gamma shape parameters (Table S5). The AtpA+B proteins both fall in the top 10 percentile of the slowest evolving genes, furthering their case for inclusion in phylogenetic analyses, considering they have likely not been transferred recently between the Nanohaloarchaea and the Haloarchaea. Furthermore, the clustering of slowly evolving genes (like the ATPases) and fast evolving genes (other genes, see Figure 4 and Table S5 for speed ranks) into the same evolutionary trajectory indicates the gene family clustering method described here may not be very sensitive to LBA artifacts. This is likely due to the analyses focusing on the correlations of individual pairwise distance matrices of each gene family.

### Analysis of Ribosomal Data

To explore the possible synergy of using multiple loci in reconstructing the history of these halophilic lineages, we constructed supermatrices of core genes and ribosomal proteins from a taxonomic sampling similar to the ATP synthase trees. We first created a ribosomal supermatrix containing 44 concatenated ribosomal proteins (RBS supermatrix). The placement of the Haloarchaea and the Nanohaloarchaea calculated from the ribosomal supermatrix (Figure S6b) resemble many previously calculated large-scale phylogenies based on RBS proteins (Andrade *et al.,* 2015; Sorokin *et al.,* 2017). In contrast to the single protein trees, the RBS tree confidently places the Nanohaloarchaea in a monophyletic clade with the rest of the DPANN members (in both the original and recoded supermatrices). The phylogeny of the 16S+23S rRNA genes recovers the Nanohaloarchaea as a sister-group to the Haloarchaea (Figure S10); however this sister-group is placed outside of the euryarchaeota. The reliability of the rRNA phylogeny is questionable, as several other groups (i.e., several DPANN members, Methanococcales, etc.) have moved from their accepted positions. These placements may reflect rRNA HGT between various Archaeal lineages (Boucher, *et al*., 2004) (Table S4), and compositional biases (in the rRNA genes) driven by adaptation to various environmental pressures.

### Analysis of Nanohaloarchaeal core genome

We also created a core genome matrix composed of 282 loci (called the Large Core supermatrix) of all genes represented in every single Nanohaloarchaeal genome considered complete. In the previously described analysis, we clustered the gene families based on the correlation between the distance matrices for these 282 gene families. From these clusters we created 3 other supermatrices which are subsets of the Large Core supermatrix, these are called the Left, Right and Center supermatrices (corresponding to the positions of the clusters on the correlated gene family dendrogram; Figure 4). Although all 282 gene families in the Large Core are represented in each Nanohaloarchaeal genome, these gene families must be analyzed for DTL conflicts and suitability in a concatenated alignment. For an example, the AtpA+B genes recover the Nanohaloarchaea and Haloarchaea as a sister-group, and falls within the Left supermatrix. However, not all large clusters of gene families (Figure 4) should fall under the assumption of vertical inheritance (one possible evolutionary trajectory). These dendrograms show differential transfer patterns of gene family clusters, and it is possible a large cluster (*i.e.,* Left or Right) could be an entire ensemble of transferred genes (another possible evolutionary trajectory). Splitting up the Large Core families into subdivisions (like the Left, Right, and Center supermatrices) may be a method to dissects common evolutionary trajectories contained in the Nanohaloarchaeal core genome.

All phylogenies calculated using the Large Core supermatrix (282 genes), and its derivatives (Left, Right and Center) recover the Nanohaloarchaea with other members of DPANN (Figure 5). In all of these supermatrices, except for the entire Large Core supermatrix, the Methanonatronarchaeia is resolved at the base of the Methanotecta group, as reported by (Aouad *et al*., 2019; Martijn *et al*., 2020). In addition, the Center and Left supermatrices recover the Haloarchaea as a sister-group to the Methanomicrobiales, similar to (Aouad *et al*., 2018). Curiously, using the Right supermatrix (Figure 5d) recovers the Haloarchaea as a sister-group to the Nanohaloarchaea. However, the Haloarchaea are relocated to outside of the euryarchaeota, and the Nanohaloarchaea are not placed outside the DPANN. This indicates that the Right cluster of genes (94 total) may contain genes frequently transferred from the Nanohaloarchaea to the Haloarchaea, representing traces of the Haloarchaea-Nanohaloarchaea symbiotic interaction. One of the pitfalls of using large concatenates is illustrated in Figure 5bc, when the genes from the Left cluster and the Right cluster are combined into a single concatenate. These clusters have clearly different phylogenetic signals (Figure 5c for the Left, Figure 5d for the Right), and forcing them on the same tree (Figure 5b) clearly leads to the one of the signals being overwhelmed by the other, with little effect on overall bootstrap support (the Right cluster is overwhelmed in this case).

The conflict of the Nanohaloarchaeal core gene families is revealed once again when the Large Core and the Left supermatrices were recoded into 4 Dayhoff groups to reduce compositional biases (Figure S7). The recoded Large Core phylogeny recovers the monophyly of all three halophilic lineages inside of the Methanotecta, with high support. However, the recoded Left core subdivision recovers the Nanohaloarchaea inside of a monophyletic DPANN. In recoded Left supermatrix, the Methanonatronarchaeia were recovered at the base of the Methanotecta. We also note that while the gene family clustering shows a better correlation for families belonging to the same cluster, the reconstructed phylogenies for individual gene families within each cluster are not identical (most individual gene families have poor phylogenetic resolution) and not identical to the phylogeny calculated from the concatenation, e.g., the AtpA and AtpB proteins are part of cluster the left cluster (Figure 5c), whose phylogeny reconstructed from the concatenation is different from the ATPase phylogeny.

In an attempt to assess the impact of long branch attraction, the DPANN sequences (excluding the Nanohaloarchaea) were removed from the entire Large Core genome supermatrix. The phylogenetic tree calculated from this new supermatrix (Figure S8) places the Nanohaloarchaea, and Haloarchaea together as sister-groups, but this sister-group has been moved out of their accepted position in the Methanotecta, and possibly out of the euryarchaeota altogether (see discussion).

### Topology Tests of Halophile Monophyly

To further investigate the possible monophyly of the three lineages, we used trees that recovered the Nanohaloarchaea, Haloarchaea, and Methanonatronarchaeia inside of the euryarchaeota as a monophyletic group (topology from Figure S7a). The explanatory power of these constrained monophyletic trees were tested against the Large and Left supermatrices using the approximately unbiased test (AU-test, Shimodaira, 2002). The topology test revealed that these constrained trees differed significantly in their likelihood landscape as compared to signals in both supermatrices, with p-values near 0. The AU-test reveals that the explanatory power of the trees with a monophyletic grouping of halophilic archaea is not compatible with the supermatrices and thus the hypothesis of halophile monophyly should be rejected under these datasets.

We also employed gene concordance factor analysis (gCF (Ané *et al*., 2007; Gadagkar *et al*., 2005)) to our Large Core supermatrix to dissect which topology each individual gene family supported. Reference trees were used to reflect conflicting evolutionary hypotheses (a tree grouping the Nanohaloarchaea and Haloarchaea together (Figure S7a) vs. a tree that placed the Nanohaloarchaea in the DPANN (Figure 5c)). Using a reference tree that was constrained to group the Nanohaloarchaea with the Haloarchaea, 16 genes were found to recover the internode that supports these groups’ monophyly. These 16 genes include highly conserved proteins such as the ATP synthase operon, ribosomal proteins, and an elongation initiation factor. In contrast, we also used an alternative reference tree that was constrained to group the Nanohaloarchaea within the DPANN superphylum and found 15 different genes in support of this hypothesis. These DPANN supporting genes include RNA polymerase subunits, FtsY, and ribosomal proteins. It is worth noting that the genes supporting the Nanohaloarchaea+Haloarchaea group consist of genes that evolve significantly slower (average rank ∼74) than those of the DPANN support set (average rank ∼93), based on the rate of evolution rankings discussed previously. We also applied the gCF method to locate the best supported placement of the Methanonatronarchaeia, and found the highest concordance was at the base of Methanotecta super class. The full table of concordant genes, as well as reference trees can be found in Figures S11,12 and Table S6.

## Discussion

### Sisterhood of Nanohaloarchaea and Haloarchaea

Analysis of the catalytic and non-catalytic subunits of the archaeal ATP synthase group the enzyme from Nanohaloarchaea as a sister-group to the Halobacterial (Haloarchaeal) subunits (Figure 2; Wang *et al.,* 2019). This strongly supported grouping is also recovered when the data are recoded to reduce compositional bias, when alignment columns containing acidic residues in both the Nanohaloarchaea and the Haloarchaea are deleted, and when the CAT-GTR model (a model that is less sensitive to compositional effects and long branch attraction artifacts) is used in phylogenetic reconstruction. None of these analyses recovered the DPANN clan, however, this may not be strong evidence against the existence of DPANN, as HGT in members of DPANN is largely unexplored in this work.

Placement of the individual DPANN groups in the ATP synthase phylogenies and the absence of an ATPase in some ectoparasitic Nanoarachaeota (Wurch *et al*., 2016) may be interpreted as evidence questioning whether the ancestor of the DPANN even possessed a functional ATP synthase. The *Nanoarchaeum equitans* genome is an example of a DPANN member which encodes the typical archaeal ATPase headgroup (3 A and 3 B subunits), although it may lack ATP synthesis activity (Mohanty *et al*., 2015). However, homology between the F (from Bacteria) and A type (from Archaea) ATPases demonstrates that these ATPases are older than the origin of Archaea (Gogarten *et al*., 1989; Gogarten & Taiz, 1992). While ATPases are known to have been horizontally transferred, the described findings suggest that the Nanoarchaeal ancestor possessed an ATPase. It is certainly possible that modern DPANN genomes replaced their ATP synthases with homologs from their hosts or from other organisms occupying the same environment.

Given the consistent support for the Nanohaloarchaea-Haloarchaea clade in the AtpA and AtpB phylogenies, it is unlikely that this finding is due to compositional bias or long branch attraction. Two conflicting hypotheses can reconcile our findings with those of previous analyses based on concatenation of several genes or on gene tree/species tree reconciliations: (1) the ATP synthase was acquired by the ancestor of the Nanohaloarchaea from a relative of the Haloarchaea or (2) the previous multi-locus analyses do not reflect evolutionary history, but are artifacts due to high substitution rates, gene transfer, and small genomes; and the Nanohaloarchaea and Haloarchaea share a common ancestor.

The recent study by Wang *et al*. (2019) includes a phylogeny derived from the entire ATPase operon in Archaea, that also recovered the sisterhood between the Nanohaloarchaea and Haloarchaea. Wang *et al*. consider horizontal transfer of the operon as explanation for this grouping, and also observe an identical operon structure in both groups, which supports the monophyly of nanohaloarchaeal and haloarchaeal ATPases. Those authors recognized clear conflicts between a DPANN supergroup and the ATPases phylogeny, and reconciled this conflict by invoking ATPase horizontal gene transfer. In correlations of Nanohaloarchaea gene families, it was revealed the ATPase genes are more closely clustered to a large set of genes; in contrast Thaumarchaeota ATPase genes form their own evolutionary cluster distant from other genes in the genome. These results can be reconciled by considering that ATPase genes were indeed transferred into Thaumarchaeota, but not in the case of Nanohaloarchaea. The gene family correlation method would be highly sensitive to single and multiple recent gene transfers, as the distance matrices analyzed for each gene family are vectorized (i.e., taxon specific information is saved and kept consistent throughout the entire clustering process, so if a single gene is transferred into only a single taxon, it will be recorded and thus affect the clustering of the entire gene family). While the clustering based on gene distance correlations does well in recovering ATPase transfer in Thaumarchaeota, the absence of detected transfer events in Nanohaloarchaea cannot be considered proof that no transfer has happened. A suspected transfer would have occurred greater than ∼1 BY ago before the deepest splits within the Nanohaloarchaea and Haloarchaea, respectively. Within-niche transfer of ATPase operons is certainly possible and is supported in the case of the Thaumarchaeota (Wang *et al.,* 2019, Figure S5) and Deinococcaceae (Lapierre *et al*., 2006); however, we are not aware of any evidence that extends this logic to the Nanohaloarchaea.

We provide evidence that ATPases do not stand out as atypical in their evolutionary history as compared to other genes found in Nanohaloarchaea (Figures 3-4, S4-5). Our interpretation of these data is that the sisterhood relationship of the Nanohaloarchaea and Haloarchaea ATPases should not be immediately discarded as resulting from HGT. The analyses presented here and by Rayman *et al.,* 2014 and Aouad *et al.,* 2018-9 suggest that Nanohaloarchaeal genomes have been shaped by a complex evolutionary history. Many gene families support inclusion of the Nanohaloarchaea into DPANN, while the aforementioned studies suggest of a placement within Haloarchaea or other euryarchaeota. The totality of support for a Nanohaloarchaea-Haloarchaea sister-group contained within our analyses include: ATPase phylogenies (and that these gene are unlikely to have been transferred (Figures 2-4, S4-5)); recoded nanohaloarchaeal Large core genome; and gCF analyses which identified several slowly evolving genes also supporting a Nanohaloarchaea-Haloarchaea sister-group relation.

Due to the observation of radically different phylogenetic signals present in the nanohaloarchaeal core, we consider an analogy between Nanohaloarchaea and Thermotoga. The Thermotoga core genome is extremely chimeric: its evolutionary history indicates genes comprising the “informational” functionality (i.e., genes involved in replication, repair, etc) are bacterial in origin, while genes that contribute to metabolism are of Archaeal or Clostridial origin (Logsdon & Faguy, 1999; Zhaxybayeva *et al*., 2009). In comparison, genes that compose the Left cluster of genes in Nanohaloarchaea are enriched for proteins that encode for translational functions, while the Right cluster is enriched for proteins that serve transcription purposes (Table S5, K= transcription, J=translation), indicating that Nanohaloarchaea are highly chimeric too.

Phylogenies calculated from concatenated datasets support the existence and monophyly of the DPANN superphylum (including the Nanohaloarchaea). When genomes from DPANN members were included, the Nanohaloarchaea were recovered as part of the DPANN group. In the absence of the other DPANN genomes, Nanohaloarchaea formed a clade with Haloarchaea (Figure S8), even after removing potential biases. However, the sister-group moved out of Methanotecta, and possibly the euryarchaeota too. As to whether this sister-group is located in the euryarchaeota depends on where one places the archaeal root. If one expects the root to be inside the euryarchaeota (Raymann *et al*., 2015), this sister-group has a possibility of falling outside the euryarchaeota, as it falls outside Methanotecta and may lead to a branch where the DPANN superphylum could attach. The observation that a monophyletic Nanohaolarchaeal-Haloarchaeal grouping is recovered from the Large core concatenation but at the base or even outside the euryarchaeota illustrates observed evolutionary relationships between archaeal classes obtained from gene family concatenations have to be interpreted with caution.

Phylogenetic reconstruction that constrained Nanohaloarchaea to group with Haloarchaea resulted in a maximum likelihood phylogeny that the AU-test (Shimodaira, 2002) evaluated as incompatible with the best tree for this Large Core genome dataset, revealing a strong phylogenetic signal, either due to shared ancestry or systematic artifact, that does contradict the sister-group relationship between Nanohaloarchaea and Haloarchaea.

Radically different placements of Nanohaloarchaea (Figure 1, red indicators) can be at least partially attributed to the taxonomic sampling of the DPANN superphylum. In instances where the Nanohaloarchaea were recovered inside the euryarchaeota (Narasingarao *et al*., 2012, Zhaxybayeva *et al*., 2013; Aouad *et al.,* 2018; Aouad *et al.,* 2019), DPANN sequences were not included in the tree. However, including a robust sampling of DPANN sequences in the alignment (Andrade *et al.,* 2015; Sorokin *et al.,* 2017; Wang *et al.,* 2019, Figure 5) generally attracts the Nanohaloarchaea into that superphylum.

The gCF analysis revealed 16 core genes in support of Nanohaloarchaea-Haloarchaea sister-group; however, 15 genes support Nanohaloarchaea inclusion in DPANN. In support of Nanohalo+Haloarchaea group are 16 genes that evolve significantly slower than those in support of the opposing hypothesis (Table S6). Previous analyses have indicated high bootstrap support for including the Nanohaloarchaea within DPANN (Sorokin *et al*., 2017; Wang *et al*., 2019). This support may reflect the strong but artifactual signal in fast evolving genes, phylogenetic signals created through gene transfers, and forcing all genes with different histories onto the same tree -- conflicting signals are likely abundant of in all concatenated marker sets. Our gCF analysis dissected the concatenation based on individual gene trees, revealing opposing phylogenetic signals present in the original concatenated dataset. It is important to supplement the sampling variance measure for the singular branch (*i.e.,* bootstrap), with a measure of variance in the overall dataset with metrics like the concordance factors. The concordance factors can reveal variance (conflict) within the multi-locus alignment datasets.

In an attempt to dissect the Large Core genome concatenation even further, we subdivided it into 3 subdivisions; the Left, Right and Center supermatrices (Figure 4). These subdivisions are based on pairwise distance matrices of each individual gene family. Phylogenies of the Right supermatrix reveals that a large ensemble of genes (94) that are a part of the Nanohaloarchaea core genome may have been transferred from the Nanohaloarchaea to Haloarchaea, as Haloarchaea moved from euryarchaeota into DPANN. Concatenations involving these genes may calculate a phylogeny with high artifact potential due to their possible transfer or unconventional evolutionary trajectory. It is worth noting again, that the AtpA+B genes fall into the Left supermatrix, even though their individual signal robustly groups the Nanohaloarchaea and Haloarchaea as sister-groups. This demonstrates that genes evolving through a similar evolutionary trajectory (Left cluster), can recover evolutionary placements and may be convoluted by concatenating gene families with disparate rates of evolution.

### Monophyly of extreme halophilic archaea

The Methanonatronarchaeia did not reveal a well-supported association with any particular Archaeal group in any of these phylogenies. In the ATP synthase-based phylogenies, homologs from three members of this group were recovered as a deeper branching euryarchaeal lineage without well supported affinity to any other euryarchaeal group. Sequences from the Methanonatronarchaeia were, however, separated by at least one well supported bipartition from other halophilic archaea grouping with non-halophilic methanogens (Figure 2, S3).

A concatenation of Nanohaloarchaeal core genes reliably placed Methanonatronarchaeia (Figure 5bc) basal to the Methanotecta super-class, as proposed by Aouad *et al.,* 2019. When using the entire Large Core genome supermatrix (Figure 5a), Methanonatronarchaeia appeared as a sister-group to Haloarchaea (BV= 88). Aouad and colleagues provided evidence for three independent adaptations to high salt environments (through the salt-in strategy) in Haloarchaea, Nanohaloarchaea, and Methanonatronarchaeia (Aouad *et al.,* 2018; Aouad *et al.,* 2019). While we consider convergent evolution events rare, independent adaptations to hypersalinity resulting from salt-in strategy pressures and revealed through shifts in protein isoelectric points (Oren, 2008) have been observed in *Salinibacter* (Bacteroidetes) and *Salinicoccus* (Firmicutes) (see Figure S13), with minimal reliance on HGT from haloarchaea (Mongodin *et al*., 2005).

Methanonatronarchaeia have been deduced to employ a salt-in strategy, using intracellular potassium ion concentrations (Sorokin *et al.,* 2017), the same adaptation present in Nanohaloarchaea and Haloarchaea. However, a proteomic analysis of theoretical isoelectric point (pI) distributions reveals a less biased distribution of pIs in these methanogens compared to other proteomes of organisms that use a salt-in strategy (Haloarchaea, Nanohaloarchaea, *etc*.) (Figure S13). This distribution of theoretical pIs in Methanonatronarchaeia resembles that found in marine archaea (Figure S13), and *Halanaerobiales*. The *Halanaerobiales* follow an experimentally confirmed salt-in strategy without an acidic proteome. Instead, they hydrolyze Glutamine (Q) and Asparagine (N) to compensate for the lack of acidic amino acids (Bardavid & Oren, 2012). However, the genome of *Acethalobium arabaticum*, a member of the *Halanaerobiales*, encodes a more acidic proteome, similar to *Salinibacter* and *Salinicoccus* (Figure S13). Methanonatronarchaeia may be a similar example of independent adaptation to hypersalinity. In some Methanonatronarchaeia the concentration of intracellular potassium did not yet have a significant impact on the distribution of pIs of encoded proteins or possibly, they may also hydrolyze their N/Q residues to make their acidic conjugates, like *Halanaerobiales*.

The AtpAB dataset robustly recovers the Nanohaloarchaea-Haloarchaea sister-group. Furthermore, we provide evidence that these genes are slow evolving (Table S5), and unlikely to have been transferred recently between the groups (Fig 3-5, S4-5). An obvious caveat is that the better resolved single gene phylogeny represents only a single gene or operon, and that its phylogeny is embedded in the net-like, reticulated genome phylogeny. Data from concatenated datasets robustly recovers the Nanohaloarchaea group within DPANN (the exception being recoded Large core phylogeny, Figure S7a). However, these datasets are rife with conflict (transferred genes, genes with differing rates of evolution; gCF and Figure 4) and forcing them on a single tree likely is inappropriate. We consider phylogenetic placement of the Nanohaloarchaea an open question. A plethora of analyses using large concatenates support inclusion of Nanohaloarchaea in DPANN (Andrade *et al*., 2015; Castelle *et al*., 2015; Rinke *et al*., 2013; Dombrowski et al., 2020), but the same can be said for the opposite (Brochier-Armanet, Forterre, & Gribaldo, 2011; Petitjean *et al*., 2014; Raymann *et al*., 2014; Williams *et al*., 2015, Aouad *et al*., 2018, Aouad *et al*., 2019). Conflict between these analyses (Figure 1) may, at least in part, be due to reliance on large concatenates and forcing disparate evolutionary signals onto the same tree. Dissecting these evolutionary signals and evaluating their suitability for such analyses could be a way forward for resolving the debate of the nanohaloarchaeal placement and the existence of DPANN.

Recently, Hikarchaeia (Martijn *et al*., 2020) were found to be more closely related to Haloarchaea (than Nanohaloarchaea) in the ATPase phylogenies. A hallmark of a salt-in strategist can be found in the Hikarchaeia’s proteomes, as they decidedly favor acidic residues found in other salt-in strategists (Figure S13mn; Oren *et al*., 2008, Paul *et al*., 2008). Results of ATPase phylogenetic analysis (Figure 2; Figure S3) also raises the possibility that both the Hikarchaeia and Haloarchaea evolved from an extreme halophilic ancestor, and that Hikarchaeia lost this adaptation after their divergence. If the ATPase phylogeny topology results from HGT or gene sharing, acquisition of the Haloarchaeal ATPase by Nanohaloarchaea predates the split between Hikarchaeia and Haloarchaea, again suggesting that extreme halophily might have been an ancestral character of the Hikarchaeia. The alternative assumption that the Hikarchaeia and Haloarchaea ancestor was not an extreme halophile would imply that transfer of the ATPase operon occurred before Haloarchaea and Nanohaloarchaea convergently adapted to hypersalinity. The inclusion of the Hikarchaeia in future phylogenetic analyses (once there is a larger sampling of this lineage) may further elucidate the genomic events that lead to hypersaline adaptation. While our analyses do not prove that Nanohaloarchaea are not part of a DPANN grouping, our findings indicate that when they are strongly supported in concatenated datasets this might be the result of an artifact, and that the phylogenies of conserved slowly evolving genes (ATPases, ribosomal proteins, and an elongation initiation factor) may better reflect the origin of the Nanohaloarchaea. In most gene families the phylogenetic signal regarding relationships between different archaeal classes is weak, and single gene phylogenies are poorly resolved. The popular solution of data set concatenation (Lapierre *et al*., 2014) to amplify a weak phylogenetic signal comes with the possibility that systematic artifacts and not a combined phylogenetic signal dominate the resulting phylogenies (Bapteste *et al*., 2008). Resolving deep divergences remains a hard problem. Due to horizontal gene transfer and phylogenetic reconstruction artifacts the placement of divergent archaeal classes into larger groups remains uncertain.

## Methods

### Sample collection, DNA extraction, and sequencing of new genomes

Two hypersaline environments in Israel were sampled for metagenomic sequences: the Dead Sea and hypersaline pools at the Mediterranean coast in Atlit. Briefly, water samples from the Dead Sea (31°30’07.2′N 35°28’37.2′E) were extracted using Niskin bottles in late July 2018. To create the enriched media, the Dead Sea water (DSW) was diluted with autoclaved double distilled water (DDW) (final ratio ⅕ [DDW/DSW]), amended with 0.1% glycerol, 1 uM KH2PO4, 1 g/L peptone (Bacto, New South Wales, Australia), 1 g/L casamino acids (Difco, Detroit, MI USA). The media was incubated at 30 °C for 42 days.

The Atlit environmental samples were collected from high salt tide-pools on the coast of Israel (32°42′37.3″N 34°56′32.0″E) in mid-October 2018. Harvesting of the microbial communities was performed by serial passage through filters (0.45um, 0.22um, 0.1um) (Merck KGaA, Darmstadt, Germany). Environmental samples (Atlit) were first prefiltered using filter paper No. 1 (11um pore size) (Munktell & Filtrak, Bärenstein, Germany). The filters were then kept in -80 °C until DNA extraction. DNA was extracted from the filters using DNeasy PowerLyzer PowerSoil kit (QIAGEN, Hilden, Germany) following the manufacturer’s protocol. For Dead Sea and Atlit samples, DNA purified from the 0.22um filters was used for library preparation (NuGen Celero enzymatic with UDI indexing). The libraries were ran on Illumina NovaSeq with SP flow cell, generating paired end reads (2x150bp).

Single amplified genomes (SAGs) were generated using fluorescence-activated cell sorting and multiple displacement amplification, as previously described (Zhaxybayeva *et al.,* 2013), from hypersaline salterns located in Santa Pola (Spain). Low coverage shotgun sequencing of SAGs was performed using Nextera library preparation and NextSeq 500 sequencers (Stepanauskas *et al*. 2017), resulting in an average of 377k, 2x150 bp reads per SAG. Although this number of reads is sub-optimal for high-quality genome reconstruction (Stepanauskas *et al*. 2017), they were sufficient to perform the specific analyses of this study. SAG generation and raw sequence generation were performed at the Bigelow Laboratory for Ocean Sciences Single Cell Genomics Center (scgc.bigelow.org).

### Sequence quality control

Raw reads obtained from single cell sequencing were trimmed and quality assured using Sickle v1.33 (Joshi & Fass, 2011) and FastQC v0.115 (Andrews, 2010). SPAdes v3.10.1 (Bankevich *et al*., 2012) was used to complete initial assemblies of single cell genomes, using option -sc. Contigs from the initial assembly were polished and bridged using the post-assembly Unicycler v0.4.7 pipeline (Wick *et al*., 2017), using normal and bold settings. Conflicts between normal and bold assemblies were investigated and reconciled in Bandage v0.8.1 (Wick *et al*., 2015). For the metagenome assembled genomes (MAGs), raw reads were trimmed using Trimmomatic-0.36 (Bolger *et al*., 2014) and quality assured using FastQC v0.10.1. SPAdes v3.11.0 was used to assemble the MAGs, using option -meta. Assembly Graphs were manually investigated using Bandage v0.8.1. Binning was conducted with MetaBat2, the bins that contained the nanohaloarchaeal MAGs were comprised of a single contig. The taxonomy and completeness of the MAGs and SAGs were checked with CheckM v1.0.7 (Parks *et al*., 2015), on default settings using a custom lineage marker developed specifically for Nanohaloarchaea (available on request). The assembled genomes were annotated with Archaeal mode Prokka v1.13.3 (Seemann, 2014). Sequences annotated as the ATP synthase alpha and beta subunits were retrieved from these genomes manually. The two MAGs in addition to ten high quality assemblies on NCBI were compiled in a library to identify the 282 core genes of the Nanohaloarchaea. Get_Homologues v03012018 (Contreras-Moreira & Vinuesa, 2013) with the COGtraingles v2.1 (Kristensen *et al*., 2010) and orthoMCL v1.4 (Li *et al*., 2003) algorithms (-t 0/1 option, -e option to exclude paralogs) were used to identify these “bona-fide” core genes used to comprise the core genome marker set.

### Whole genome distance analysis

Average nucleotide identity (ANI) was calculated using a slight modification of the JSpecies method (Richter & Rosselló-Móra, 2009). Genomes were divided into 1,020 nt fragments and used as the query for pairwise BLAST searches. A 70% identity and 70% coverage cutoff was implemented in a manner akin to the global ANI (gANI) filtering method (Varghese *et al*., 2015). The filtered BLASTN (Camacho *et al*., 2009) searches were also used to calculate a modified gANI and alignment fraction (AF), which were used to construct a phylogenetic tree as per the tANI method (Gosselin *et al*., 2020). The entire method and standalone script can be found at: https://github.com/SeanGosselin/tANI_Matrix.git.

### Assembly of datasets

169 high quality genomes spanning the Archaea domain were collected through NCBI’s ftp site, and were supplemented with the seven newly assembled Nanohaloarchaea genomes (Table S1). AtpA and AtpB protein sequences were found in these genomes and gathered with BLASTP v2.7.1, using default parameters. Similarly, protein sequences of 282 Nanohaloarchaea core proteins and forty-four ribosomal proteins were found and gathered from these genomes using TBLASTN using default parameters. Sequences hits from each protein were categorized into their own respective files and aligned with Mafft-linsi (Katoh & Standley, 2013). Alignments were trimmed by BMGE (Criscuolo & Gribaldo, 2010). Each alignment file of the core and ribosomal protein dataset was concatenated using FASconCAT-G (Kück & Longo, 2014) to generate supermatrices and the associated nexus partition files. A description of the core supermatrices is available in the supplemental material (Table S5).

The core supermatrices and the ATPase dataset were recoded into Dayhoff groups (4 and 6) based on functional classes of amino acids, using PhyloBayes v4.1 (Lartillot *et al*., 2009). We also manually curated alternative alignments which had removed alignment columns if they contained an Aspartate or Glutamate (D/E) residue that was conserved in the Nanohaloarchaea, Haloarchaea, and Methanonatronarchaeia, to minimize compositional attraction in the ATPase dataset.

### Phylogenetic Estimation

IQTREE v1.6.9 (Nguyen *et al*., 2015) was used to calculate maximum likelihood phylogenies for all alignments and supermatrices. The best site homogeneous models were used for the estimation as determined by the Bayesian Information Criterion using ModelFinder (Kalyaanamoorthy *et al*., 2017), for single gene phylogenies and guide tree calculation. The ATPase and concatenated alignments (also recoded versions) were also analyzed by the LG+C60 (Le *et al*., 2008) mixture model, all trees reported in the main text and supplemental text were calculated using this model. Bayesian inference of Dayhoff recoded ATPase alignments were conducted within PhyloBayes v4.1(Lartillot *et al*., 2009; Quang *et al*.,2008) using the CAT+GTR +G4 model in two independent chains for each alignment. These chains ran until convergence (maxdiff < 0.25), >400,000 trees sampled, with a burn-in of the first 10% of the trees, to calculate a majority rule consensus tree. All trees in this paper were drawn and editorialized with Figtree v1.4.3 (Rambaut, 2016). The approximately unbiased tests were also carried out in IQTREE (parameters: -zb 10000, -n 0) with multi-treefiles that contained phylogenies from the hypotheses of interest (Nanohaloarchaea in DPANN or as a sister-group to Haloarchaea) and 1000 bootstrap trees from opposing hypotheses (either from ATPases or core genome dataset). gCF analyses of the large core supermatrix was carried out in the IQTREEv1.7.17 beta.

### Clustering of Gene Families

Individual alignments of encoded proteins were gathered for two marker sets, in 3 separate clustering analyses: the 282 core gene families (which focused on the Nanohaloarchaea), and the Archaea 122 (Parks *et al*., 2018) marker set (which was compiled for the Thaumarchaeota and the Nanohaloarchaea, separately), in addition to the ATPase genes. For each taxon of interest, a sampling of genomes from the suspected transfer partners for each taxon was also included (i.e., for the Nanohaloarchaeal analysis Haloarchaeal genomes were also included; for the Thaumarchaeota genomes from Micrarchaeota and Thermoplasmatales were included (based on Wang *et al.,* 2019)). For the Thaumarchaeota, no distinction was made between those species that have received their ATPases via HGT and those that have not, both types of genomes were included. Phylogenies were calculated for all the alignments in IQTREE (using the best model determined by BIC), and pairwise distance matrices (taxon vs taxon) were generated. These pairwise distance matrices contain distances from pairwise sequence comparisons, calculated with maximum likelihood based on model parameter estimates from an initial tree for each protein alignment. Using pairwise distance matrices built from sequence comparisons avoids relying on reconstructing phylogenetic trees where the placement of our groups of interest are already uncertain. This correlation of pairwise distance matrices of gene families is based on co-evolution implementations found in (Gueudré, *et al*., 2016, Rangel *et al.,* 2020) These pairwise distance matrices were regressed (in the sklearn Python module) against all other distance matrices in each respective dataset (*i.e.,* each gene *vs* the 281 other genes, or 1 gene *vs* 121 genes). The ability of one distance matrix to predict the values of another distance matrix was defined by 1-r^2^, and a summary pairwise distance matrix (gene family vs gene family) was calculated. These values were the basis of agglomerative clustering implemented in Agnes (cluster v2.1.0), which computes all pairwise dissimilarities between gene families and considers the average of a pair the distance on the clustering diagram, and generated the clustered gene families in Figure S4-5. The full implementation and instructions of this method can be found in the gene_fam_dist.ipynb file in https://github.com/Gogarten-Lab-Team/NanoH_GBE_2020.

## Acknowledgments and Funding

This work was supported by grants from the Binational Science Foundation [BSF 2013061 to U.G., J.P.G., and R.T.P.]; the National Science Foundation [NSF/MCB 1716046 to J.P.G., R.T.P., and U.G.] within the BSF-NSF joint research program; and the National Aeronautics and Space Administration exobiology [NNX15AM09G and 80NSSC18K1533 to R.T.P.]. We thank the staff of the Bigelow Laboratory for Ocean Sciences’ Single Cell Genomics Center for the generation of single cell genomic data. We specifically thank Ramunas Stepanauskas for helpful critical discussions and overseeing the single cell sequencing. We also thank Dr. Thiberio Rangel for his insightful input and the code base for the gene family clustering analyses, and Dr. Jan Gogarten for his input with the nMDS plots.

## Author Contributions

The project was conceived by RTP, JPG, and UG. Sampling, sequencing, and genome reconstruction of Dead Sea samples were conducted by UN and UG. Single cell samples were collected by RTP, and assemblies performed by ASL and YF. Phylogenies were calculated by YF, whole genome distance by SG. All authors contributed to writing and editing the manuscript.

## Data Availability

The newly assembled Nanohaloarchaeal genomes and accompanying information have been deposited into GenBank under BioProject PRJNA587522 for the SAGs, and PRJNA588232 for the MAGs. Alignments and raw newick treefiles are archived in https://github.com/Gogarten-Lab-Team/NanoH_GBE_2020.

## Supplementary Materials Table of Contents

### Supplementary Figures (supp_files.pdf)

**S1**. tANI Distance Matrix to classify new Nanohaloarchaea.

**S2**. Classification of new Nanohaloarchaea genomes with tANI distances.

**S3**. Selected phylogenies of the Archaeal ATP synthase subunits.

**S4**. Clustering diagram of clustered core gene families in the Nanohaloarchaea.

**S5**. Diagram of clustered gene families in the Nanohaloarchaea, using the Archaeal 122 marker set.

**S6**. Diagram of clustered gene families in the Thaumarchaeota.

**S7**. Phylogenetic trees calculated from a supermatrix of ribosomal proteins.

**S8**. Phylogeny of recoded core supermatrices.

**S9**. Phylogeny of the Large core supermatrix, with DPANN sequences removed.

**S10**. Phylogenies of the Large core supermatrix with fast evolving sites stripped from the alignment.

**S11**. Phylogeny of the 16S+23S rRNA concatenate.

**S12**. Constrained reference tree used to test the monophyly of the halophiles in gCF analyses.

**S13**. Constrained reference tree used to test the hypothesis of the Nanohaloarchaea falling in DPANN in gCF analyses.

**S14**. Protein theoretical isoelectric point distribution in selected proteomes.

**S15.** Figure S4 with tip labels.

**S16.** Clustering of gene families from Thaumarchaeota subsets.

**Table S6**. Concordant gene trees with each evolutionary hypothesis.

### Supplementary Tables (Supplementary_tables1-5.xlsx)

**Table S1.** Genome files and associated statistics used in this study.

**Table S2.** Assembled genome statistics.

**Table S3.** Assembled genome gene content.

**Table S4.** Summary of phylogenetic analyses.

**Table S5.** Description of large core genome markers.

