## Supplementary material for "The evolutionary origins of extreme halophilic Archaeal lineages": Supp_materials: supp_figs.pdf

**Note**: all figures are zoomable

| | Hfx. gibbonsii AR6 | Hfx. mediterranei | Hfx. volcanii | Hrr. californiensis | Hrr. ezzemulense | Hrr. lacusprofundi | Hrr. lacusprofundi HLS1 | Hrr. trapanicum | Mca. arvory |
|---|---|---|---|---|---|---|---|---|---|
| Hfx. gibbonsii AR6 | 0.00 | 1.17 | 0.45 | 2.63 | 2.64 | 2.76 | 2.77 | 2.62 | 6.44 |
| Hfx. mediterranei | 1.14 | 0.00 | 1.18 | 3.30 | 3.25 | 3.23 | 3.25 | 3.25 | 7.16 |
| Hfx. volcanii | 0.47 | 1.23 | 0.00 | 2.50 | 2.46 | 2.58 | 2.61 | 2.61 | 6.78 |
| Hrr. californiensis | 2.54 | 3.18 | 2.49 | 0.00 | 0.82 | 1.37 | 1.40 | 0.80 | 7.45 |
| Hrr. ezzemulense | 2.57 | 3.14 | 2.50 | 0.81 | 0.00 | 1.37 | 1.39 | 0.89 | 7.43 |
| Hrr. lacusprofundi | 2.66 | 3.06 | 2.48 | 1.33 | 1.31 | 0.00 | 0.18 | 1.29 | 6.38 |
| Hrr. lacusprofundi HLS1 | 2.60 | 3.11 | 2.57 | 1.28 | 1.27 | 0.07 | 0.00 | 1.22 | 7.66 |
| Hrr. trapanicum | 2.43 | 3.00 | 2.35 | 0.62 | 0.70 | 1.16 | 1.17 | 0.00 | 7.04 |
| Mca. arvory | 6.04 | 6.14 | 6.13 | 6.46 | 6.59 | 6.11 | 6.11 | 6.63 | 0.00 |
| Mca. conradii HZ254g | 6.69 | 6.97 | 6.97 | 6.76 | 6.77 | 6.77 | 6.77 | 6.77 | 3.74 |
| Mca. paludicola | 6.59 | 6.81 | 6.80 | 6.62 | 6.79 | 6.47 | 6.47 | 6.61 | 3.13 |
| Msc. barkeri 3 | 6.96 | 6.98 | 6.97 | 7.03 | 7.02 | 6.97 | 6.98 | 7.00 | 6.22 |
| Msc. barkeri MS | 6.91 | 6.90 | 6.91 | 6.88 | 6.92 | 6.90 | 6.92 | 6.91 | 6.27 |
| Msc. thermophila | 6.53 | 6.55 | 6.53 | 6.61 | 6.59 | 6.55 | 6.55 | 6.58 | 5.82 |
| nano AB578 D14 | 13.00 | 13.00 | 13.00 | 13.00 | 13.00 | 7.39 | 7.49 | 7.11 | 13.00 |
| nano B1 Br10 U2g1 | 13.00 | 13.00 | 13.00 | 13.00 | 13.00 | 13.00 | 13.00 | 13.00 | 13.00 |
| nano B1 Br10 U2g19 | 13.00 | 13.00 | 13.00 | 13.00 | 13.00 | 13.00 | 13.00 | 13.00 | 13.00 |
| nano B1 Br10 U2g21 | 13.00 | 13.00 | 13.00 | 13.00 | 13.00 | 13.00 | 13.00 | 13.00 | 13.00 |
| nano B1 Br10 u2g29 | 13.00 | 13.00 | 13.00 | 13.00 | 13.00 | 13.00 | 13.00 | 13.00 | 13.00 |
| nano PL Br10 u2g27 | 13.00 | 13.00 | 13.00 | 13.00 | 13.00 | 13.00 | 13.00 | 13.00 | 13.00 |
| nano PL Br10 u2g29 | 13.00 | 13.00 | 13.00 | 13.00 | 13.00 | 13.00 | 13.00 | 13.00 | 13.00 |
| nano PL br10 u2g5 | 2.72 | 3.18 | 2.73 | 1.40 | 1.42 | 1.30 | 1.30 | 1.42 | 13.00 |
| nano Pl Br10 U2g16 | 13.00 | 13.00 | 13.00 | 13.00 | 13.00 | 13.00 | 13.00 | 13.00 | 13.00 |
| nano QH8446 | 13.00 | 13.00 | 13.00 | 13.00 | 13.00 | 13.00 | 13.00 | 13.00 | 13.00 |
| nano SG9 | 13.00 | 13.00 | 13.00 | 13.00 | 13.00 | 13.00 | 13.00 | 13.00 | 13.00 |
| nano SW104410 | 13.00 | 13.00 | 13.00 | 13.00 | 13.00 | 13.00 | 13.00 | 13.00 | 13.00 |
| nano SW4439 | 13.00 | 13.00 | 13.00 | 13.00 | 13.00 | 13.00 | 13.00 | 13.00 | 13.00 |
| nano SW7431 | 13.00 | 13.00 | 13.00 | 13.00 | 13.00 | 13.00 | 13.00 | 13.00 | 13.00 |
| nano SW7467 | 13.00 | 13.00 | 13.00 | 13.00 | 13.00 | 13.00 | 13.00 | 13.00 | 13.00 |
| MAG AT 22 | 13.00 | 13.00 | 13.00 | 13.00 | 13.00 | 13.00 | 13.00 | 13.00 | 13.00 |
| MAG M3 22 | 13.00 | 13.00 | 13.00 | 13.00 | 13.00 | 13.00 | 13.00 | 13.00 | 13.00 |
| Nanosalina J07AB43 | 7.72 | 7.70 | 7.72 | 13.00 | 13.00 | 13.00 | 13.00 | 13.00 | 13.00 |
| Nanosalinarum J07AB56 | 7.55 | 7.55 | 7.55 | 7.37 | 6.36 | 7.72 | 7.72 | 7.37 | 13.00 |
| SAG M04 | 13.00 | 13.00 | 13.00 | 13.00 | 13.00 | 13.00 | 13.00 | 13.00 | 13.00 |
| SAG M06 | 13.00 | 13.00 | 13.00 | 13.00 | 13.00 | 13.00 | 13.00 | 13.00 | 13.00 |
| SAG M21 | 13.00 | 8.73 | 13.00 | 6.57 | 6.40 | 6.83 | 8.13 | 7.01 | 13.00 |

| | Mca. conradii HZ254g | Mca. paludicola | Msc. barkeri 3 | Msc. barkeri MS | Msc. thermophila | nano AB578 D14 | nano B1 Br10 U2g1 | nano B1 Br10 U2g19 | nano B1 Br10 U2g21 |
|---|---|---|---|---|---|---|---|---|---|
| Hfx. gibbonsii AR6 | 6.92 | 6.44 | 7.13 | 7.11 | 7.11 | 13.00 | 13.00 | 13.00 | 13.00 |
| Hfx. mediterranei | 7.46 | 6.95 | 7.20 | 7.20 | 7.20 | 13.00 | 13.00 | 13.00 | 13.00 |
| Hfx. volcanii | 7.51 | 6.80 | 7.18 | 7.17 | 7.17 | 13.00 | 13.00 | 13.00 | 13.00 |
| Hrr. californiensis | 7.55 | 7.46 | 7.72 | 7.70 | 7.87 | 13.00 | 13.00 | 13.00 | 13.00 |
| Hrr. ezzemulense | 7.14 | 7.19 | 7.11 | 7.11 | 7.11 | 8.76 | 13.00 | 13.00 | 13.00 |
| Hrr. lacusprofundi | 6.82 | 6.65 | 6.62 | 6.62 | 6.63 | 7.96 | 13.00 | 13.00 | 13.00 |
| Hrr. lacusprofundi HLS1 | 7.81 | 8.48 | 7.72 | 7.72 | 7.66 | 8.99 | 13.00 | 13.00 | 13.00 |
| Hrr. trapanicum | 7.07 | 6.67 | 7.00 | 6.81 | 6.82 | 8.45 | 13.00 | 13.00 | 13.00 |
| Mca. arvory | 3.93 | 3.21 | 5.86 | 5.86 | 5.78 | 13.00 | 13.00 | 13.00 | 13.00 |
| Mca. conradii HZ254g | 0.00 | 2.10 | 6.42 | 6.62 | 6.42 | 13.00 | 13.00 | 13.00 | 13.00 |
| Mca. paludicola | 2.32 | 0.00 | 6.46 | 6.45 | 6.33 | 13.00 | 13.00 | 13.00 | 13.00 |
| Msc. barkeri 3 | 6.62 | 6.42 | 0.00 | 0.95 | 1.75 | 13.00 | 13.00 | 13.00 | 13.00 |
| Msc. barkeri MS | 6.49 | 6.28 | 0.94 | 0.00 | 1.74 | 13.00 | 13.00 | 13.00 | 13.00 |
| Msc. thermophila | 6.10 | 5.98 | 1.41 | 1.37 | 0.00 | 13.00 | 13.00 | 13.00 | 13.00 |
| nano AB578 D14 | 13.00 | 13.00 | 13.00 | 13.00 | 13.00 | 0.00 | 7.30 | 7.06 | 13.00 |
| nano B1 Br10 U2g1 | 13.00 | 13.00 | 13.00 | 13.00 | 13.00 | 6.97 | 0.00 | 2.06 | 5.33 |
| nano B1 Br10 U2g19 | 13.00 | 13.00 | 13.00 | 13.00 | 13.00 | 6.59 | 1.92 | 0.00 | 4.71 |
| nano B1 Br10 U2g21 | 13.00 | 13.00 | 13.00 | 13.00 | 13.00 | 13.00 | 5.46 | 5.09 | 0.00 |
| nano B1 Br10 u2g29 | 13.00 | 13.00 | 13.00 | 13.00 | 13.00 | 13.00 | 4.46 | 4.64 | 3.97 |
| nano PL Br10 u2g27 | 13.00 | 13.00 | 13.00 | 6.74 | 13.00 | 6.43 | 4.01 | 4.18 | 2.04 |
| nano PL Br10 u2g29 | 13.00 | 13.00 | 13.00 | 13.00 | 13.00 | 6.66 | 0.27 | 2.10 | 6.43 |
| nano PL br10 u2g5 | 13.00 | 13.00 | 13.00 | 13.00 | 13.00 | 13.00 | 13.00 | 13.00 | 13.00 |
| nano Pl Br10 U2g16 | 13.00 | 13.00 | 13.00 | 13.00 | 13.00 | 13.00 | 4.98 | 5.16 | 5.52 |
| nano QH8446 | 13.00 | 13.00 | 13.00 | 13.00 | 13.00 | 13.00 | 5.79 | 4.58 | 6.79 |
| nano SG9 | 13.00 | 13.00 | 13.00 | 13.00 | 13.00 | 6.50 | 5.43 | 4.93 | 6.30 |
| nano SW104410 | 13.00 | 13.00 | 13.00 | 13.00 | 13.00 | 13.00 | 3.89 | 4.21 | 5.62 |
| nano SW4439 | 13.00 | 13.00 | 13.00 | 13.00 | 13.00 | 6.27 | 4.76 | 4.29 | 6.11 |
| nano SW7431 | 13.00 | 13.00 | 13.00 | 13.00 | 13.00 | 6.39 | 4.85 | 4.88 | 7.30 |
| nano SW7467 | 13.00 | 13.00 | 13.00 | 13.00 | 13.00 | 13.00 | 4.94 | 5.20 | 5.65 |
| MAG AT 22 | 13.00 | 13.00 | 13.00 | 13.00 | 13.00 | 7.16 | 4.44 | 3.97 | 6.69 |
| MAG M3 22 | 13.00 | 13.00 | 13.00 | 13.00 | 13.00 | 6.28 | 5.14 | 4.51 | 5.86 |
| Nanosalina J07AB43 | 13.00 | 13.00 | 13.00 | 13.00 | 13.00 | 6.73 | 7.53 | 5.66 | 6.79 |
| Nanosalinarum J07AB56 | 13.00 | 13.00 | 13.00 | 13.00 | 13.00 | 2.32 | 13.00 | 7.50 | 13.00 |
| SAG M04 | 13.00 | 13.00 | 13.00 | 13.00 | 13.00 | 4.33 | 5.10 | 4.92 | 6.72 |
| SAG M06 | 13.00 | 13.00 | 7.76 | 7.76 | 7.76 | 5.47 | 6.15 | 5.70 | 7.32 |
| SAG M21 | 13.00 | 13.00 | 13.00 | 13.00 | 13.00 | 1.46 | 13.00 | 7.24 | 13.00 |

| | nano B1 Br10 u2g29 | nano PL Br10 u2g27 | nano PL Br10 u2g29 | nano PL br10 u2g5 | nano Pl Br10 U2g16 | nano QH8446 | nano SG9 | nano SW104410 | nano SW4439 |
|---|---|---|---|---|---|---|---|---|---|
| Hfx. gibbonsii AR6 | 13.00 | 13.00 | 13.00 | 3.72 | 13.00 | 13.00 | 13.00 | 13.00 | 13.00 |
| Hfx. mediterranei | 13.00 | 13.00 | 13.00 | 4.29 | 13.00 | 13.00 | 13.00 | 13.00 | 13.00 |
| Hfx. volcanii | 8.68 | 13.00 | 13.00 | 3.73 | 13.00 | 13.00 | 13.00 | 13.00 | 13.00 |
| Hrr. californiensis | 13.00 | 13.00 | 13.00 | 2.38 | 13.00 | 13.00 | 13.00 | 13.00 | 8.58 |
| Hrr. ezzemulense | 13.00 | 13.00 | 13.00 | 2.36 | 13.00 | 13.00 | 7.60 | 13.00 | 13.00 |
| Hrr. lacusprofundi | 13.00 | 13.00 | 13.00 | 2.23 | 13.00 | 13.00 | 13.00 | 13.00 | 13.00 |
| Hrr. lacusprofundi HLS1 | 13.00 | 13.00 | 13.00 | 2.12 | 13.00 | 13.00 | 13.00 | 13.00 | 13.00 |
| Hrr. trapanicum | 13.00 | 13.00 | 13.00 | 2.20 | 13.00 | 13.00 | 8.71 | 13.00 | 13.00 |
| Mca. arvory | 13.00 | 13.00 | 13.00 | 13.00 | 13.00 | 13.00 | 13.00 | 13.00 | 13.00 |
| Mca. conradii HZ254g | 13.00 | 13.00 | 13.00 | 13.00 | 13.00 | 13.00 | 13.00 | 13.00 | 13.00 |
| Mca. paludicola | 13.00 | 13.00 | 13.00 | 8.49 | 13.00 | 13.00 | 13.00 | 13.00 | 13.00 |
| Msc. barkeri 3 | 13.00 | 8.94 | 13.00 | 13.00 | 13.00 | 13.00 | 13.00 | 13.00 | 13.00 |
| Msc. barkeri MS | 13.00 | 13.00 | 13.00 | 13.00 | 13.00 | 13.00 | 9.06 | 13.00 | 13.00 |
| Msc. thermophila | 13.00 | 13.00 | 13.00 | 13.00 | 13.00 | 13.00 | 13.00 | 13.00 | 13.00 |
| nano AB578 D14 | 13.00 | 6.44 | 6.61 | 13.00 | 13.00 | 13.00 | 6.21 | 13.00 | 6.46 |
| nano B1 Br10 U2g1 | 4.94 | 4.62 | 0.23 | 13.00 | 4.90 | 13.00 | 4.65 | 4.68 | 4.81 |
| nano B1 Br10 U2g19 | 4.59 | 4.24 | 1.96 | 13.00 | 5.11 | 5.09 | 4.36 | 4.75 | 4.56 |
| nano B1 Br10 U2g21 | 4.21 | 2.53 | 5.45 | 13.00 | 5.70 | 7.23 | 5.68 | 7.16 | 7.15 |
| nano B1 Br10 u2g29 | 0.00 | 3.60 | 4.42 | 13.00 | 5.27 | 5.56 | 4.61 | 5.89 | 6.62 |
| nano PL Br10 u2g27 | 3.74 | 0.00 | 3.94 | 13.00 | 5.97 | 6.07 | 4.44 | 5.31 | 4.93 |
| nano PL Br10 u2g29 | 5.17 | 4.49 | 0.00 | 13.00 | 5.02 | 5.96 | 4.53 | 4.68 | 4.97 |
| nano PL br10 u2g5 | 13.00 | 13.00 | 13.00 | 0.00 | 13.00 | 13.00 | 13.00 | 13.00 | 13.00 |
| nano Pl Br10 U2g16 | 5.69 | 6.79 | 5.15 | 13.00 | 0.00 | 6.79 | 5.23 | 5.76 | 5.45 |
| nano QH8446 | 5.68 | 5.23 | 6.35 | 13.00 | 6.67 | 0.00 | 1.42 | 1.66 | 1.28 |
| nano SG9 | 5.14 | 5.18 | 5.04 | 13.00 | 6.00 | 2.50 | 0.00 | 5.78 | 4.77 |
| nano SW104410 | 5.62 | 4.57 | 3.95 | 13.00 | 6.26 | 1.41 | 4.91 | 0.00 | 0.46 |
| nano SW4439 | 6.78 | 4.81 | 4.88 | 13.00 | 6.25 | 1.76 | 4.05 | 1.19 | 0.00 |
| nano SW7431 | 6.49 | 5.21 | 4.65 | 13.00 | 5.60 | 1.99 | 4.40 | 1.45 | 0.91 |
| nano SW7467 | 4.99 | 6.67 | 5.14 | 13.00 | 5.44 | 2.18 | 0.36 | 6.17 | 6.09 |
| MAG AT 22 | 5.58 | 5.32 | 4.32 | 13.00 | 4.95 | 5.84 | 4.48 | 4.77 | 4.45 |
| MAG M3 22 | 4.97 | 4.43 | 5.15 | 13.00 | 5.32 | 5.63 | 3.71 | 5.01 | 5.33 |
| Nanosalina J07AB43 | 6.16 | 6.28 | 5.97 | 13.00 | 7.44 | 7.42 | 5.10 | 6.41 | 5.69 |
| Nanosalinarum J07AB56 | 13.00 | 6.35 | 6.66 | 13.00 | 7.62 | 13.00 | 5.78 | 7.56 | 6.63 |
| SAG M04 | 5.45 | 5.67 | 5.11 | 13.00 | 13.00 | 13.00 | 5.07 | 13.00 | 7.59 |
| SAG M06 | 5.59 | 5.55 | 5.59 | 13.00 | 6.24 | 6.63 | 4.79 | 6.09 | 5.56 |
| SAG M21 | 13.00 | 7.26 | 7.00 | 13.00 | 7.76 | 7.53 | 5.69 | 13.00 | 5.64 |

| | nano SW7431 | nano SW7467 | MAG AT 22 | MAG M3 22 | Nanosalina J07AB43 | Nanosalinarum J07AB56 | SAG M04 | SAG M06 | SAG M21 |
|---|---|---|---|---|---|---|---|---|---|
| Hfx. gibbonsii AR6 | 13.00 | 13.00 | 13.00 | 13.00 | 13.00 | 8.03 | 13.00 | 13.00 | 8.67 |
| Hfx. mediterranei | 13.00 | 13.00 | 13.00 | 13.00 | 8.82 | 8.08 | 13.00 | 8.80 | 10.00 |
| Hfx. volcanii | 13.00 | 13.00 | 13.00 | 13.00 | 13.00 | 8.83 | 13.00 | 13.00 | 13.00 |
| Hrr. californiensis | 13.00 | 13.00 | 13.00 | 13.00 | 13.00 | 8.41 | 13.00 | 13.00 | 8.07 |
| Hrr. ezzemulense | 13.00 | 13.00 | 13.00 | 13.00 | 13.00 | 7.26 | 13.00 | 13.00 | 7.14 |
| Hrr. lacusprofundi | 13.00 | 13.00 | 13.00 | 13.00 | 13.00 | 8.58 | 13.00 | 13.00 | 6.80 |
| Hrr. lacusprofundi HLS1 | 13.00 | 13.00 | 13.00 | 13.00 | 13.00 | 8.39 | 13.00 | 13.00 | 7.74 |
| Hrr. trapanicum | 13.00 | 13.00 | 13.00 | 13.00 | 13.00 | 8.24 | 13.00 | 13.00 | 7.48 |
| Mca. arvory | 13.00 | 13.00 | 13.00 | 13.00 | 13.00 | 13.00 | 13.00 | 13.00 | 13.00 |
| Mca. conradii HZ254g | 13.00 | 13.00 | 13.00 | 13.00 | 13.00 | 13.00 | 13.00 | 13.00 | 13.00 |
| Mca. paludicola | 13.00 | 13.00 | 13.00 | 13.00 | 13.00 | 13.00 | 13.00 | 13.00 | 13.00 |
| Msc. barkeri 3 | 13.00 | 13.00 | 13.00 | 13.00 | 8.83 | 13.00 | 13.00 | 9.34 | 13.00 |
| Msc. barkeri MS | 13.00 | 13.00 | 13.00 | 13.00 | 8.11 | 13.00 | 13.00 | 8.11 | 13.00 |
| Msc. thermophila | 13.00 | 13.00 | 13.00 | 13.00 | 8.57 | 13.00 | 13.00 | 9.16 | 13.00 |
| nano AB578 D14 | 6.28 | 13.00 | 6.55 | 6.03 | 6.22 | 1.95 | 4.91 | 5.57 | 1.31 |
| nano B1 Br10 U2g1 | 4.32 | 5.30 | 3.98 | 4.66 | 6.44 | 13.00 | 5.57 | 6.22 | 13.00 |
| nano B1 Br10 U2g19 | 4.22 | 5.86 | 3.56 | 3.81 | 5.03 | 6.59 | 5.32 | 5.34 | 6.58 |
| nano B1 Br10 U2g21 | 6.40 | 7.25 | 5.71 | 5.88 | 6.46 | 13.00 | 7.16 | 13.00 | 13.00 |
| nano B1 Br10 u2g29 | 5.06 | 5.95 | 5.37 | 4.74 | 5.21 | 13.00 | 5.98 | 5.19 | 13.00 |
| nano PL Br10 u2g27 | 4.74 | 13.00 | 4.66 | 3.89 | 5.04 | 5.47 | 5.78 | 5.04 | 5.52 |
| nano PL Br10 u2g29 | 4.40 | 5.42 | 3.92 | 4.69 | 5.85 | 6.31 | 6.22 | 5.45 | 6.09 |
| nano PL br10 u2g5 | 13.00 | 13.00 | 13.00 | 13.00 | 13.00 | 13.00 | 13.00 | 13.00 | 13.00 |
| nano Pl Br10 U2g16 | 4.75 | 6.93 | 4.48 | 5.01 | 6.80 | 13.00 | 13.00 | 13.00 | 13.00 |
| nano QH8446 | 1.12 | 1.98 | 4.47 | 4.54 | 5.41 | 13.00 | 13.00 | 5.88 | 6.61 |
| nano SG9 | 4.65 | 1.21 | 4.70 | 3.80 | 5.12 | 5.94 | 6.05 | 5.54 | 5.73 |
| nano SW104410 | 0.35 | 13.00 | 3.55 | 3.99 | 5.07 | 6.19 | 13.00 | 5.71 | 6.20 |
| nano SW4439 | 0.52 | 5.83 | 4.26 | 4.65 | 4.70 | 5.81 | 13.00 | 5.43 | 4.99 |
| nano SW7431 | 0.00 | 13.00 | 4.57 | 4.34 | 5.71 | 5.74 | 6.92 | 5.59 | 5.05 |
| nano SW7467 | 5.48 | 0.00 | 4.63 | 4.01 | 5.23 | 13.00 | 13.00 | 5.96 | 13.00 |
| MAG AT 22 | 4.52 | 5.40 | 0.00 | 4.38 | 5.06 | 6.72 | 6.60 | 5.28 | 6.73 |
| MAG M3 22 | 4.33 | 5.17 | 4.46 | 0.00 | 4.40 | 6.03 | 5.67 | 5.40 | 5.86 |
| Nanosalina J07AB43 | 5.53 | 6.34 | 5.51 | 4.94 | 0.00 | 6.76 | 1.76 | 1.15 | 6.51 |
| Nanosalinarum J07AB56 | 6.24 | 13.00 | 7.58 | 5.81 | 6.64 | 0.00 | 7.29 | 6.85 | 2.05 |
| SAG M04 | 5.77 | 6.02 | 5.43 | 4.59 | 0.67 | 7.53 | 0.00 | 0.61 | 7.55 |
| SAG M06 | 5.01 | 6.48 | 5.23 | 5.14 | 0.82 | 6.34 | 1.41 | 0.00 | 6.23 |
| SAG M21 | 5.25 | 7.43 | 7.25 | 6.00 | 6.85 | 1.81 | 8.45 | 6.51 | 0.00 |

Fig S1. **tANI distance matrix used to calculate Fig S2, and to delineate taxonomy.** Red squares indicate a distance within the threshold of the two genomes representing the same species (d < 0.315), green squares are suggestive of the same genus (d < 3.4). Distances were calculated by the perl script tANI_beta.pl. The distance is calculated as d = -log(AF*ANI), where ANI is the average nucleotide identity, and AF is the alignment fraction as described (Gosselin *et al.* 2020).

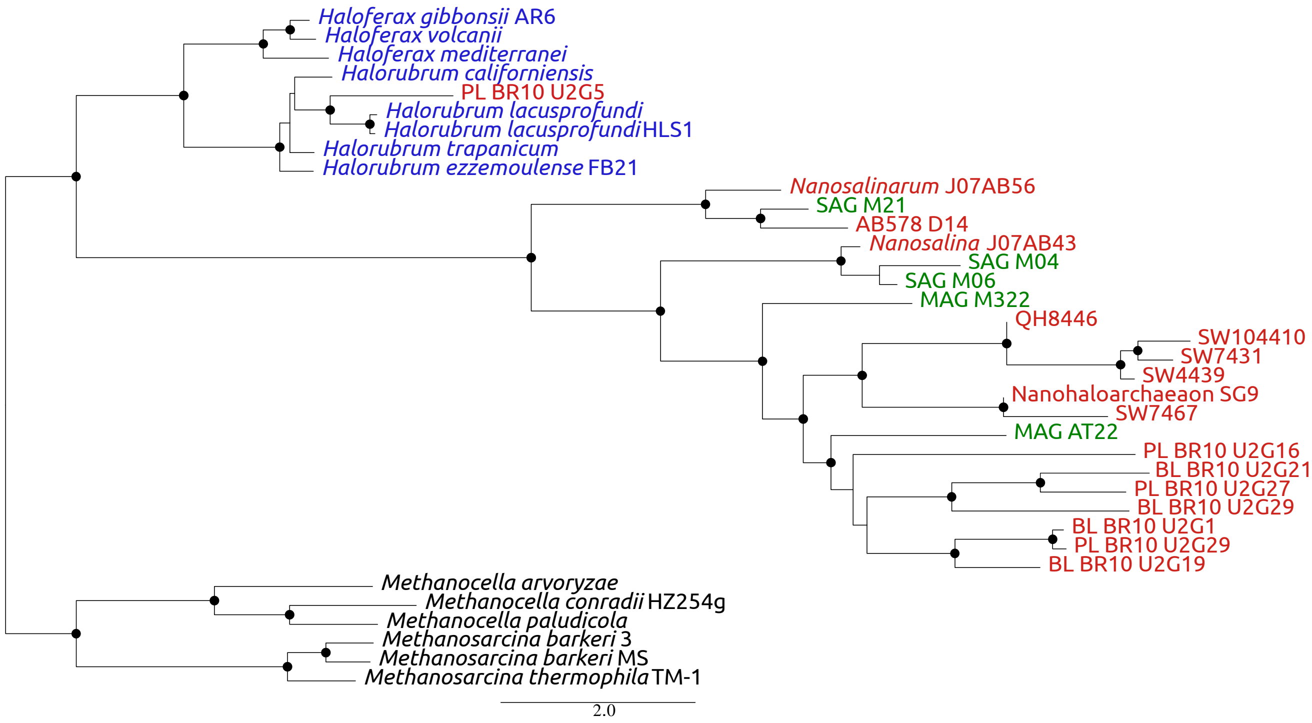


Fig S2. **Classification of new Nanohaloarchaea genomes with tANI distances.** This tree was calculated using a whole genome ANI metric that takes into account the alignment fraction of matches, and corrects distances to correct for saturation. The strains prefixed nano were retrieved from NCBI, while MAG (metagenome assembled genome) and SAG (single amplified genome) designations indicate in-house assembly. Bootstrap values were calculated from pairwise distances using sampled genomes as query. Nodes with greater than 80% support are represented with a circle at the node. Strain nano_PL-Br-U2g5 was described as a *Nanohaloarchaeon*, but groups with high support among *Halorubrum spp.* The tree should be considered un-rooted. Those taxa with green coloring indicates the genomes generated from this study.

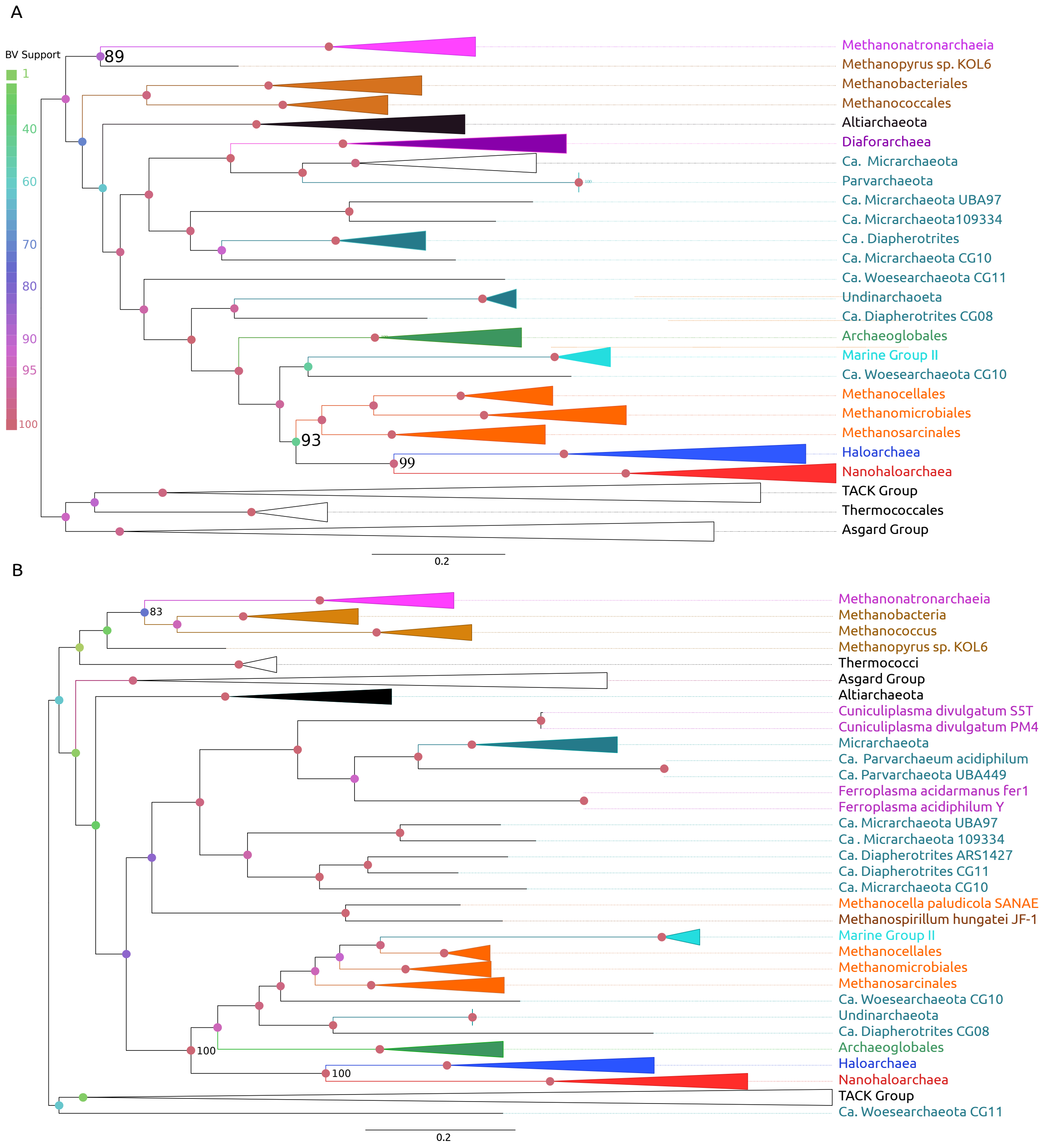


Fig S3. **Selected phylogenies of the Archaeal ATP synthase subunits**. Several chosen taxa were collapsed into higher taxonomic ranks. Branch supports are non-parametric bootstraps, most relevant supports are displayed with numerical value, otherwise all supports BV are represented by the coloring of circles at the nodes. Important taxa including the halophilic lineages and DPANN (teal) sequences have been colored; *Nanohaloarchaea* (red), *Haloarchaea* (blue), *Methanonatronarchaeia* (purple), Methanotecta methanogens (orange), Methanomada (brown), and the Hikarchaeia (magenta). Both trees were calculated with the LG+C60 model. A) Calculated from an alignment of the AtpA protein. B) Calculated from an alignment of AtpB. Note the branches breaking the DPANNs apart are sometimes highly supported; while the Haloarchaea and Nanohaloarchaea are still recovered inside of the euryarchaeota. DPANN placement seems erratic and may be indicative of HGTs.

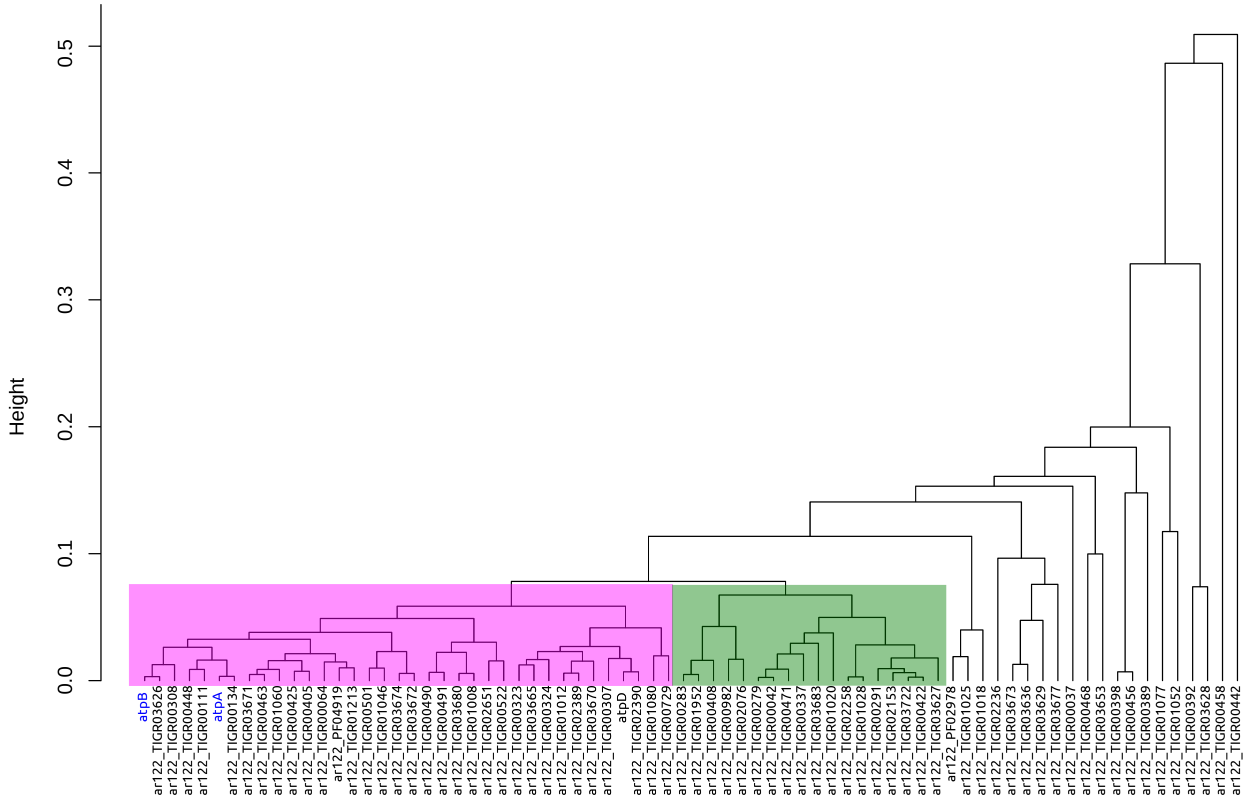


Fig S4. **Diagram of clustered gene families in the Nanohaloarchaea**. Gene family alignments from the Archaea_122 marker set in the Nanohaloarchaea clustered based on the correlation between pairwise distance matrices (gene family vs gene family). The blue tip labels indicates where the AtpA+B genes fall, with respect to other gene families. Those gene families enclosed by the purple rectangle are considered "clustered" and share evolutionary trajectories with other members of the cluster. The unenclosed branches likely contain genes that have been transferred into the Nanohaloarchaea, from divergent organisms (likely from donors that were not included in this analysis). Gene families with less than three significant homologs and with unreliable alignments were omitted from the clustering. This clustering diagram is different from Fig. S4, as the genes included are from the Archaeal 122 marker set (Parks *et al.*, 2018) and is meant to represent a reliable set of genes to compare the gene family correlations between the Nanohaloarchaea and the Thaumarchaeota.

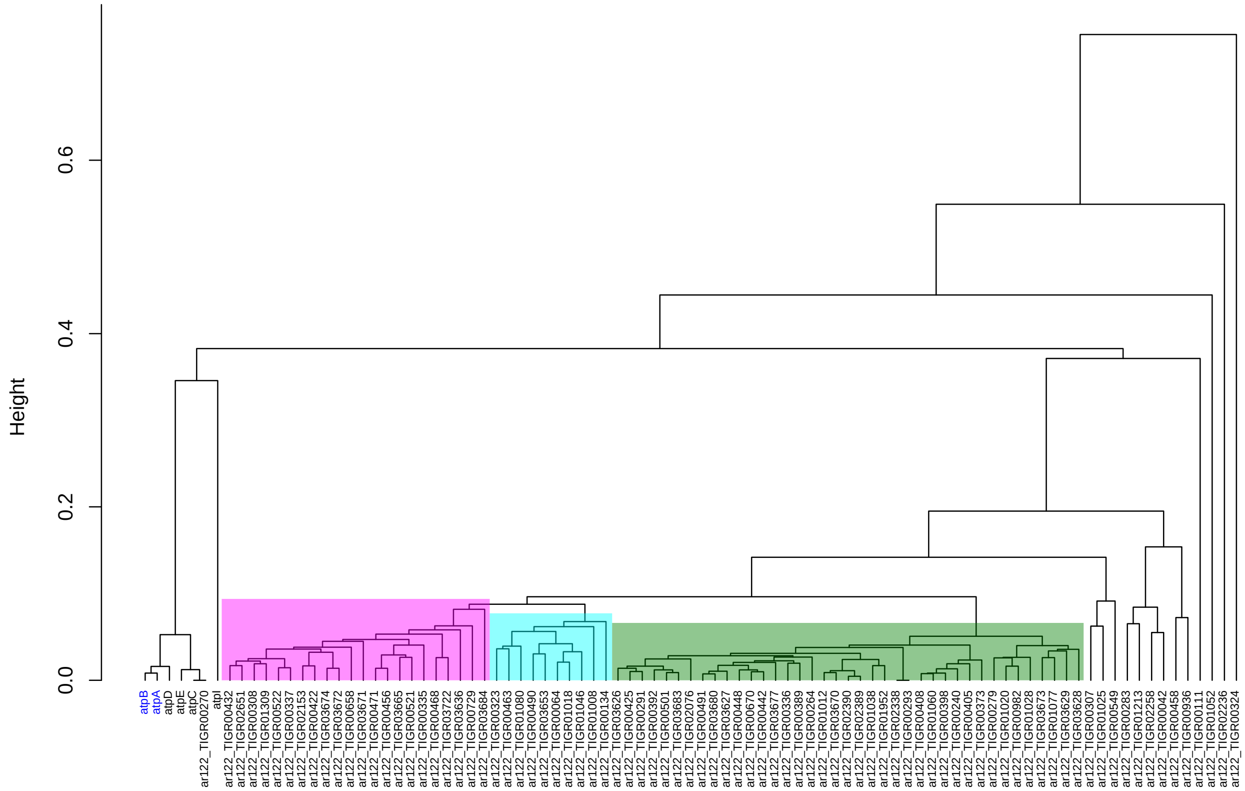


Fig S5. **Diagram of clustered gene families in the Thaumarchaeota**. Gene family alignments from the Archaea_122 marker set in the Thaumarchaeota clustered based on the correlatin between pairwise distance matrices(gene family vs gene family). The blue tip labels indicate where the AtpA+B genes fall, with respect to other gene families. Those gene families enclosed with rectangles are considered “clustered” and have similar evolutionary trajectories with other members of the cluster. The gene families on long branches have likely been transferred into the Thaumarchaeota or follow an unconventional evolutionary trajectory. Gene families with less than three significant homologs were omitted from the clustering. Note the AtpA+B genes fall outside of the main clustered groups and supports the idea that they have been transferred within the group, from a common evolutionary source.

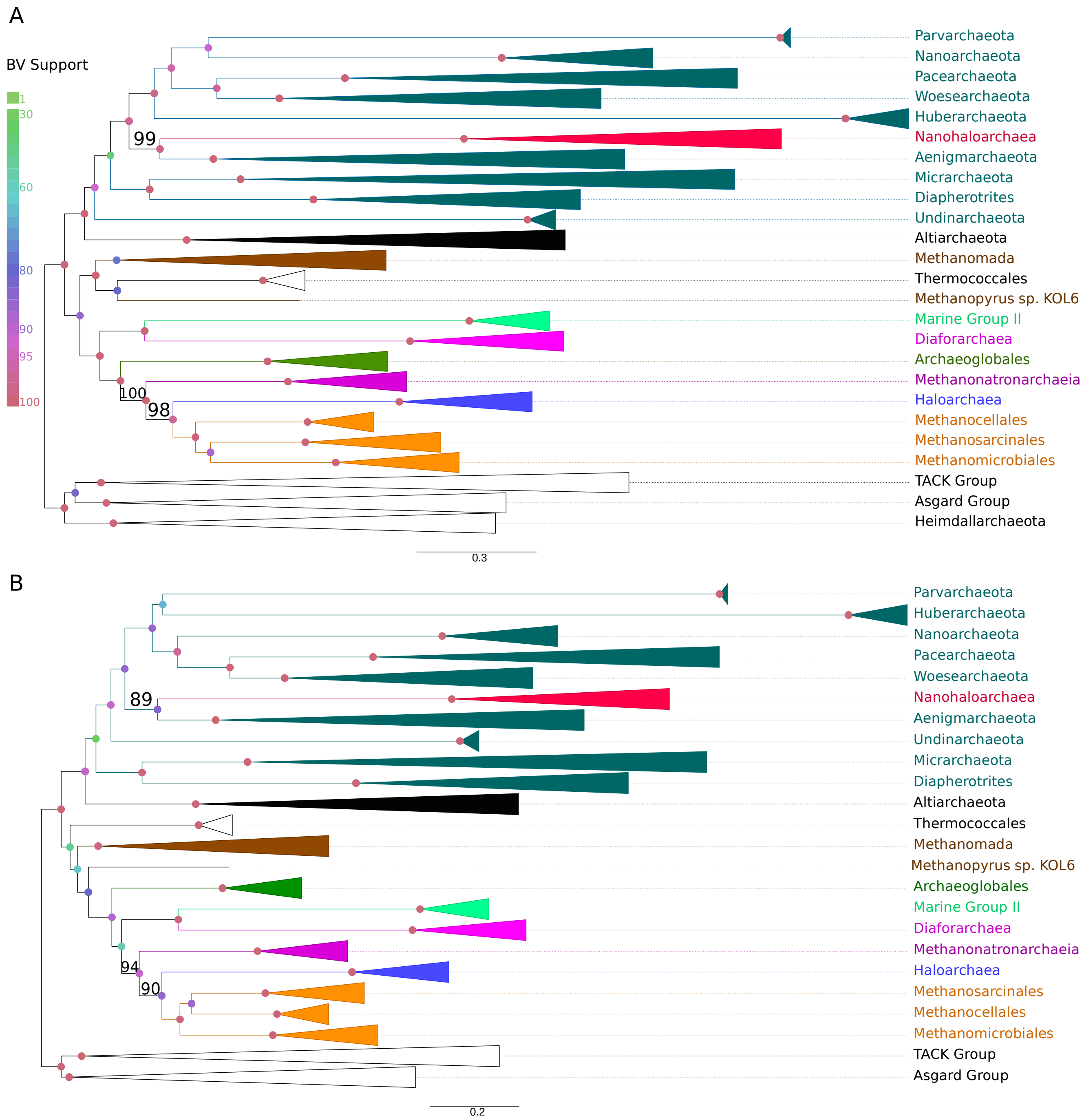


Fig S6. **Phylogenetic trees reconstructed from a supermatrix of ribosomal proteins (44 proteins).** The coloring scheme is the same as in Fig. S3, both trees were calculated using the LG+C60 models. A) Maximum likelihood phylogeny of the RBS supermatrix. The Methanonatronarchaeia are basal to the Haloarchaea, while the Nanohaloarchaea fall in the DPANN with high support. B) Built from the same ribosomal supermatrix recoded into 4 Dayhoff Groups, the placement of the halophiles are the same as in Fig S7a.

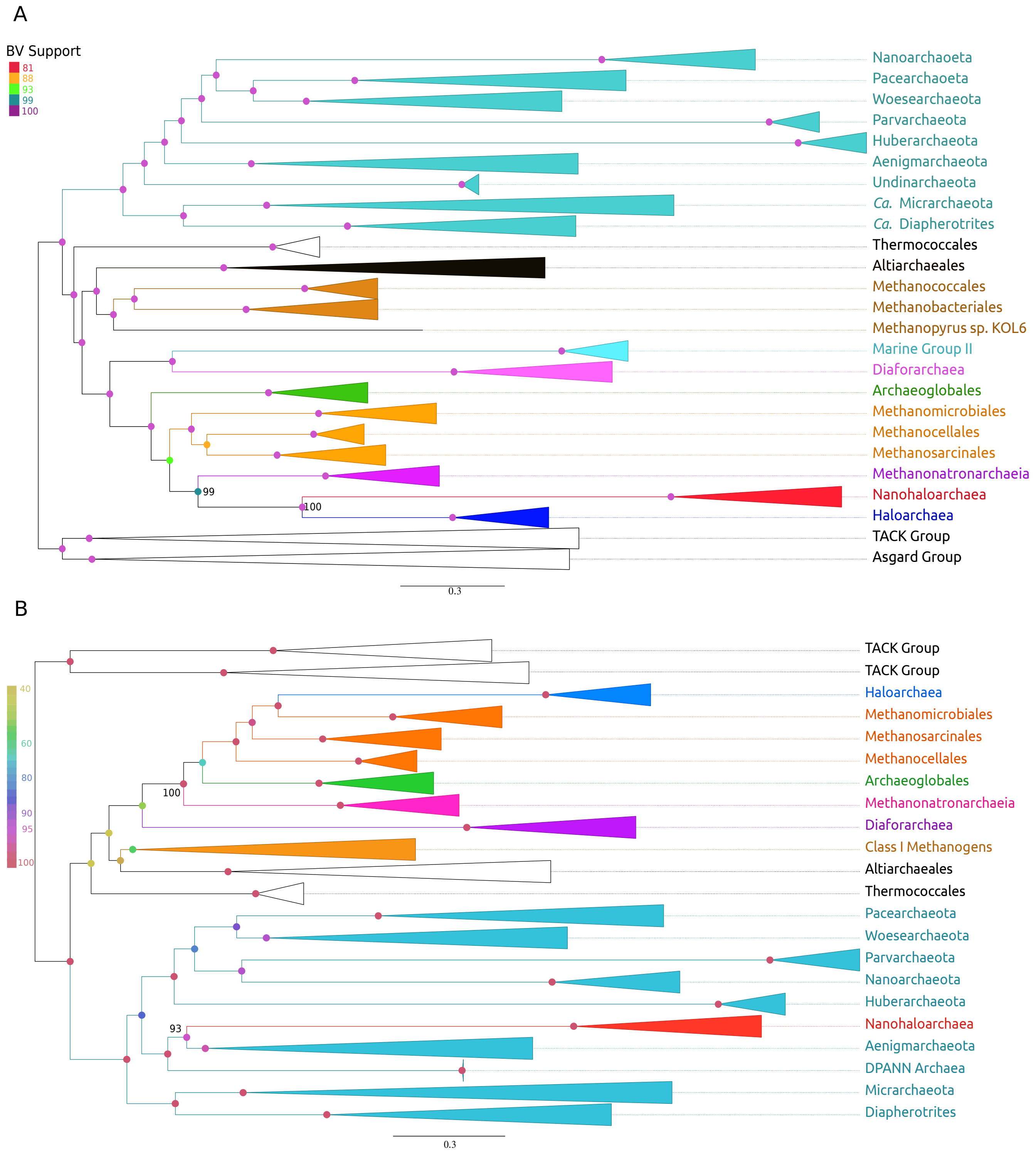


Fig S7. **Phylogeney of recoded core supermatrices.** These supermatrices contain the core genes of the Nanohaloarchaea, and were recoded into 4 Dayhoff groups and evaluated with the LG+C60 mixture model. A) ML phylogeny of the recoded 282 core genes in the Nanohaloarchaea. The Nanohaloarchaea and Haloarchaea group together with high support. The Methanonatronarchaeia are placed at the base of the Nanohaloarchaea-Haloarchaea sister group, with high support. B) ML phylogeny of the recoded Left cluster of genes. The Nanohaloarchaea fall in a monophyletic DPANN, while the Methanonatronarchaeia are placed at the base of the Methanotecta.

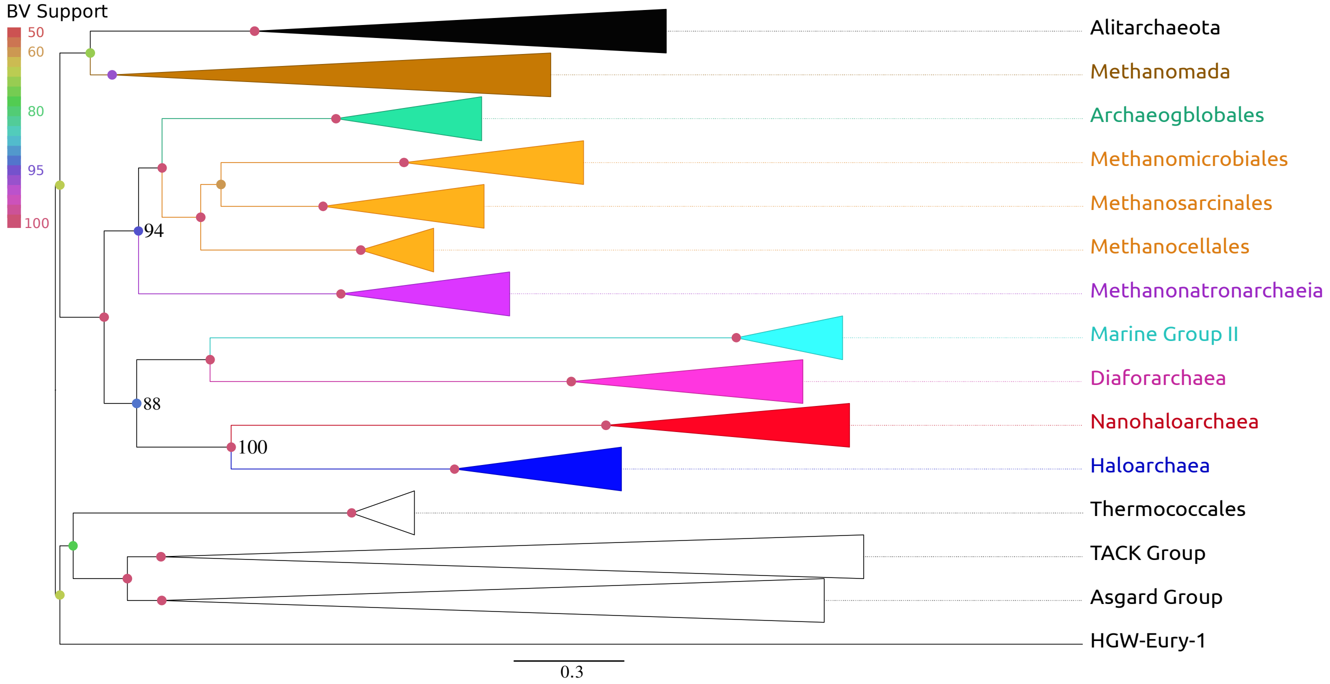


Fig S8. **Phylogeney of the Large Core supermatrix, with DPANN sequences removed.** Built from the same Large Core supermatrix without the DPANN members (using the LG+C60 mixture model), the Nanohaloarchaea group with the Haloarchaea, but this sister group has moved out of its accepted position with the Methanotecta methanogens (orange). The taxonomy of Eury. archeon HGW-Euryarchaeota-1 is unclear, its has been reported to be a euryarchaeote on NCBI taxonomy, but is also often seen as a DPANN.

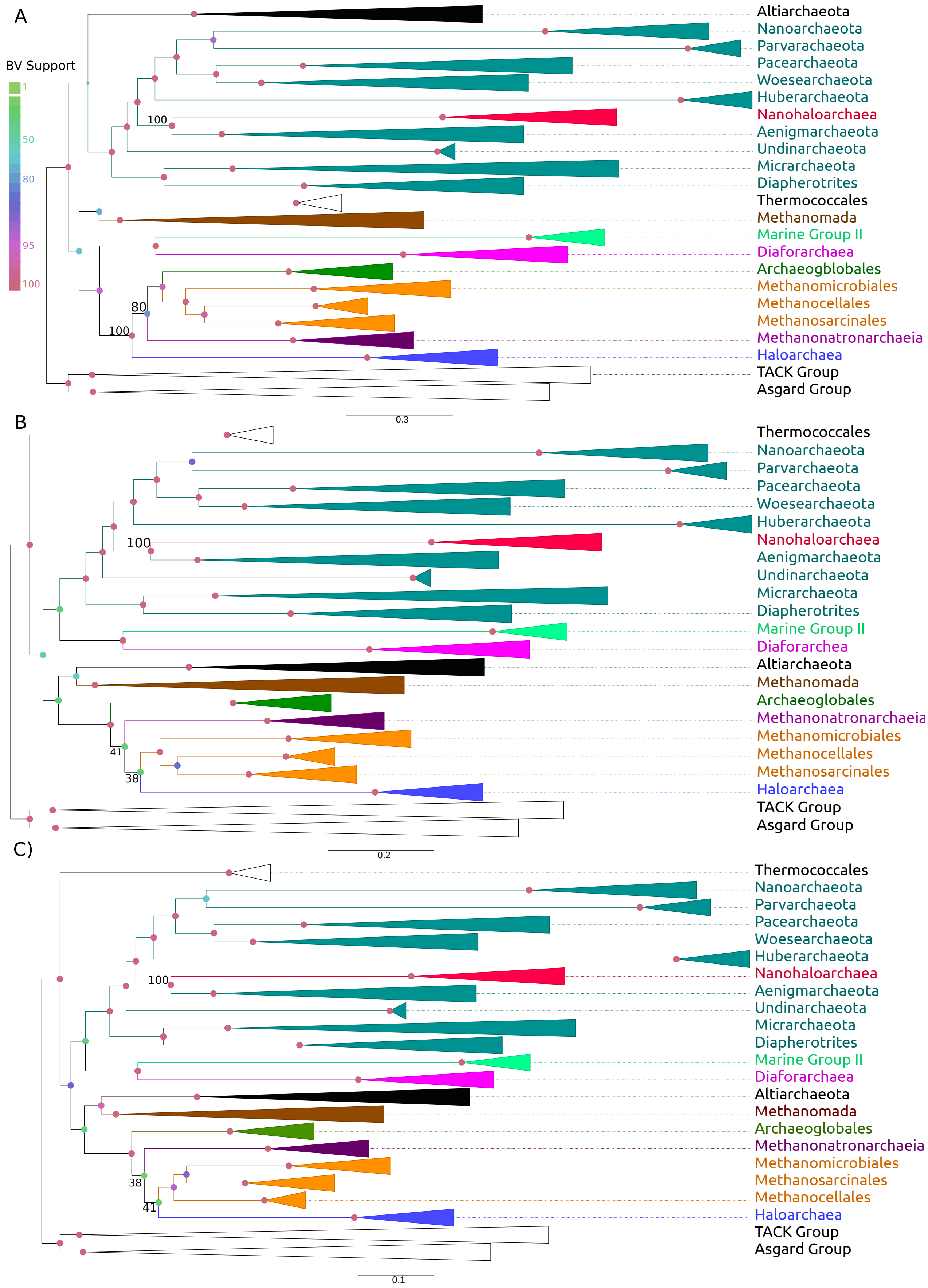
A
BV Support
1
50
80
95
100
Altiarchaeota
Nanoarchaeota
Parvarachaeota
Pacearchaeota
Woesearchaeota
Huberarchaeota
Nanohaloarchaea
Aenigmarchaeota
Undinarchaeota
Micrarchaeota
Diapherotrites
Thermococcales
Methanomada
Marine Group II
Diaforarchaea
Archaeogblobales
Methanomicrobiales
Methanocellales
Methanosarcinales
Methanonatronarchaeia
Haloarchaea
TACK Group
Asgard Group
100
80
100
0.3
B
Thermococcales
Nanoarchaeota
Parvarchaeota
Pacearchaeota
Woesearchaeota
Huberarchaeota
Nanohaloarchaea
Aenigmarchaeota
Undinarchaeota
Micrarchaeota
Diapherotrites
Marine Group II
Diaforarchea
Altiarchaeota
Methanomada
Archaeoglobales
Methanonatronarchaeia
Methanomicrobiales
Methanocellales
Methanosarcinales
Haloarchaea
TACK Group
Asgard Group
100
41
38
0.2
C)
Thermococcales
Nanoarchaeota
Parvarchaeota
Pacearchaeota
Woesearchaeota
Huberarchaeota
Nanohaloarchaea
Aenigmarchaeota
Undinarchaeota
Micrarchaeota
Diapherotrites
Marine Group II
Diaforarchaea
Altiarchaeota
Methanomada
Archaeoglobales
Methanonatronarchaeia
Methanomicrobiales
Methanosarcinales
Methanocellales
Haloarchaea
TACK Group
Asgard Group
100
38
41
0.1

Fig S9. **Phylogenies of the Large core supermatrix with fast evolving sites stripped from the alignment.** Fast evolving sites from the alignment matrix were found and categorized by using -wsr and -wsl (in IQTree) to obtain site rates and site log-likelihoods. Those sites found to be fast evolving were removed using site_stripper.pl. All of the trees were calculated using the LG+c60 model. A) Large core supermatrix where 95% of sites were retained (5% of fast evolving sites removed). B) Large core supermatrix where 80% of sites were retained. C) Large core supermatrix where 60% of sites were retained. The Nanohaloarchaea were placed in a monophyletic DPANN in all of these phylogenies, while the placement of Methanonatronarchaeia varied.

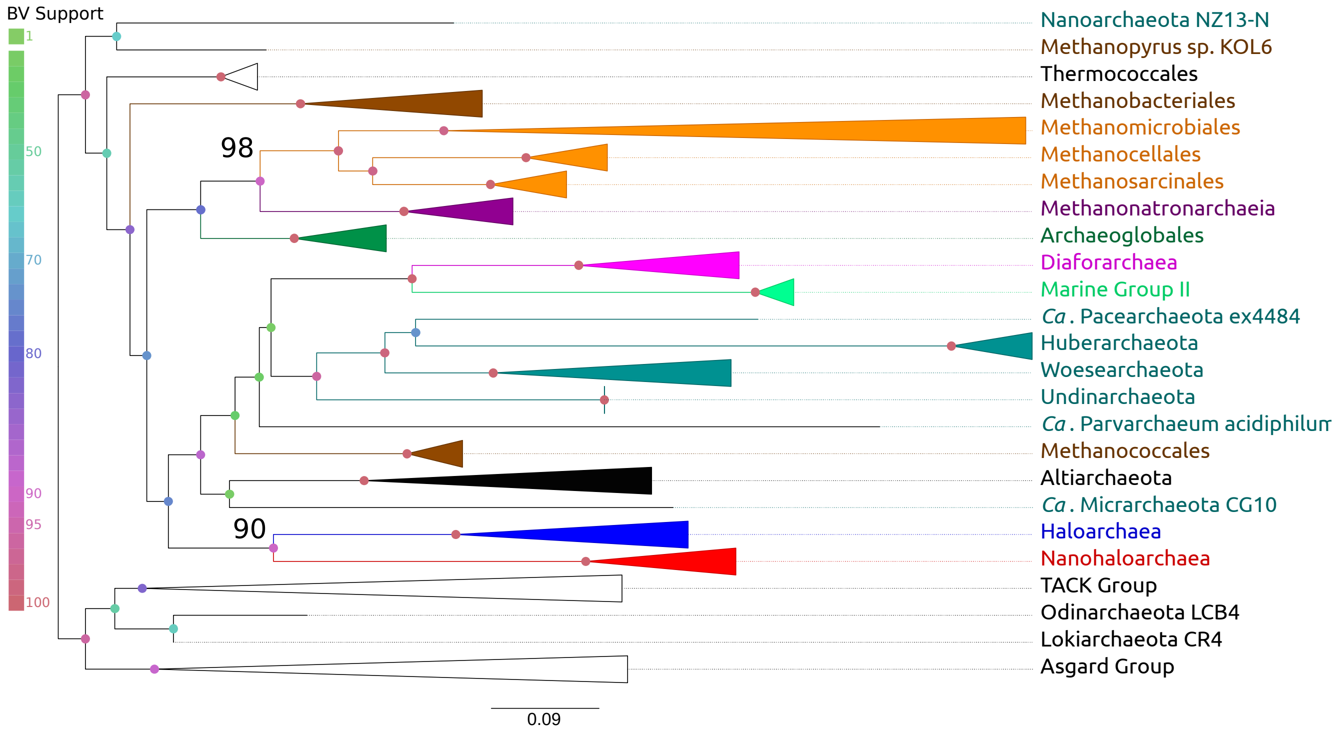


Fig S10. **Phylogeny of the 16S + 23S rRNA concatenate.** Phylogeny of concatenated ribsosomal rRNA, calculated with GTR+G4 model in IQTree, hypervariable sites were masked by 10%. The reliability of this tree is hard to assess as groups and species fall out of their expected positions, or may be indicative of HGT or homogenization. The Nanohaloarchaea-Haloarchaea sister group is recovered, but falls outside of the euryarchaeota. DPANN groups also are scattered among the tree.

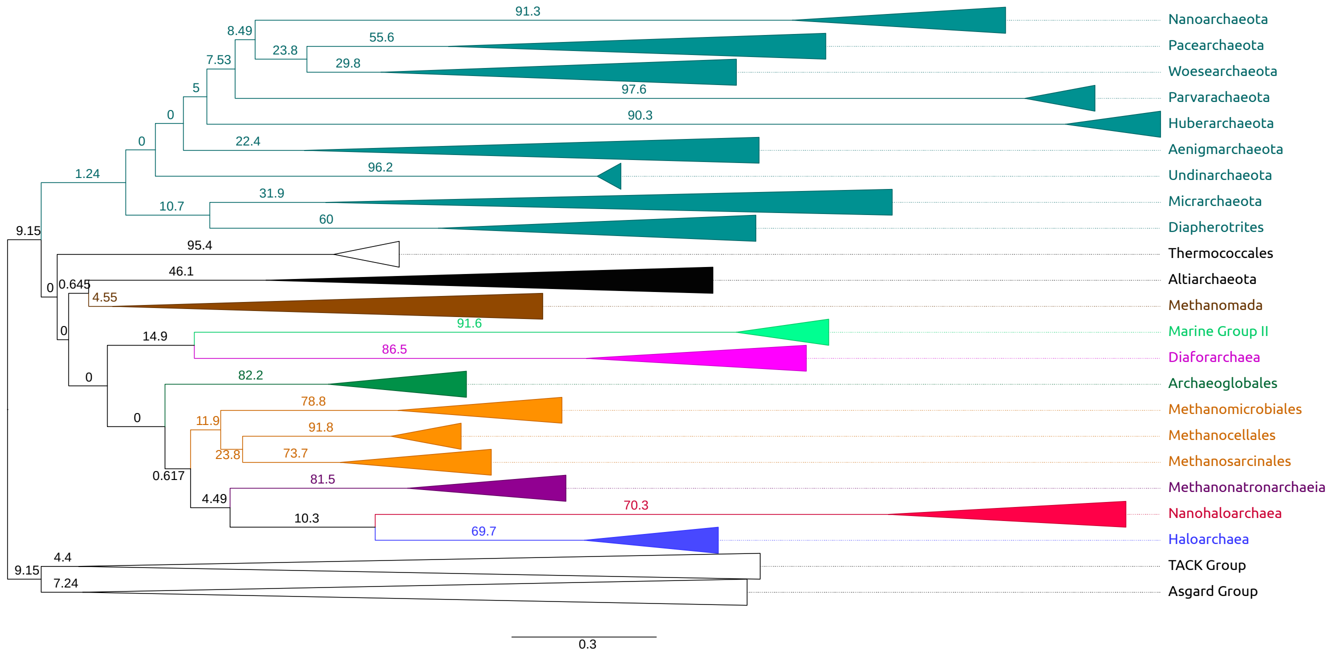


S11. **Constrained reference tree used to test the monophyly of the halophiles in gCF analyses.** The reference phylogeny used to test the hypothesis that all three halophilic lineages group together in the euryarchaeota, as a reference tree to map concordant gene families. This reference tree is the same as the tree in Fig S8a (recoded large core supermatrix), where the monophyly of the groups were recovered. Branch supports indicate the percentage of "decisive" gene trees (out of a possible maximum of 282 gene trees), that recover exactly the quartet at each node. A decisive gene is any possible gene tree that can possibly contain the branch in question (from a starting total of 282 genes). The grouping of the Nanohaloarchaea and the Haloarchaea was recovered by 16 genes or 10.3% of total decisive trees (155). Each gene family phylogeny was calculated by using the best model (determined by BIC) for each family.

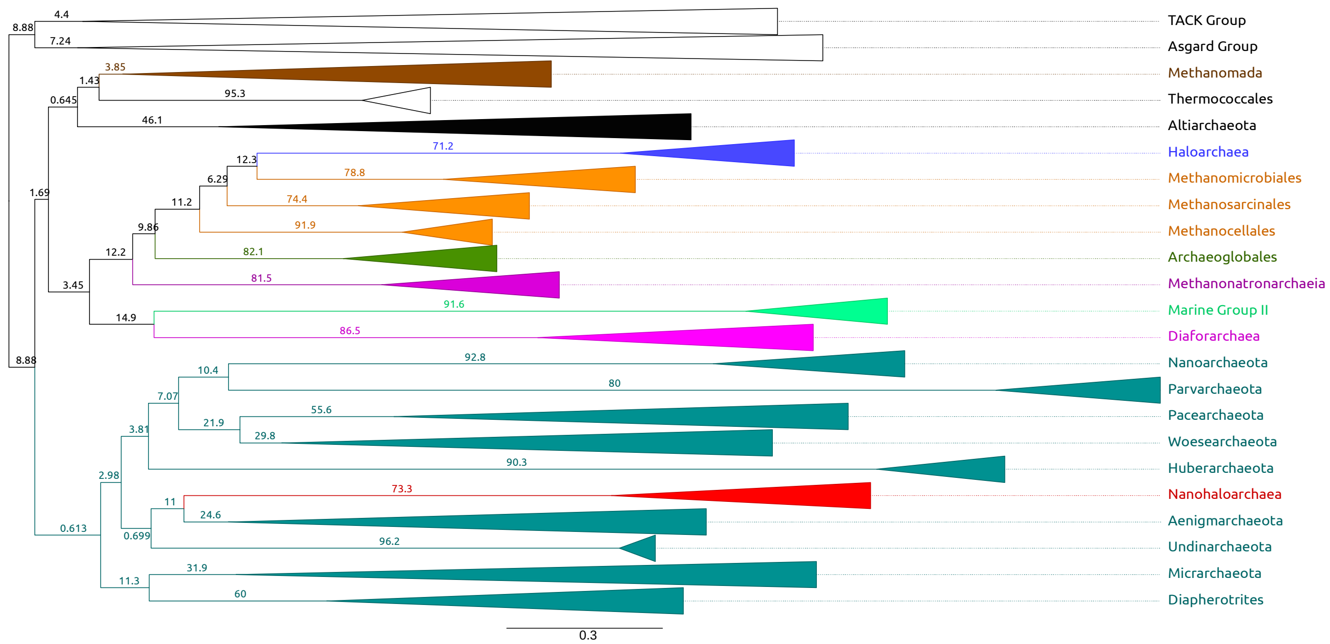


Fig S12. **Constrained reference tree used to test the hypothesis of the Nanohaloarchaea falling in the DPANN in gCF analyses.** This was used as a reference tree to map concordant gene families. The reference tree is the same as the tree calculated using the Left core supermatrix (Fig 4c), where the Nanohaloarchaea were placed in the DPANN, and the Methanonatronarchaeia were recovered at the base of the Methanotecta. Branch supports indicate the percentage of "decisive" gene trees, that recover exactly the quartet at each node. A decisive gene is any possible gene tree that can possibly contain the branch in question (from a starting total of 282 genes). The grouping of the Nanohaloarchaea in the DPANN, and as a sister group to the Aenigmarchaeota is supported by 15 genes or 11% of 127 decisive genes. The placement of the Methanonatronarchaeia at the base of the Methanotecta is supported by 15 genes or 12.2% of 131 decisive genes. Each gene family phylogeny was calculated by using the best model (determined by BIC) for each family.

Table S6. **Concordant gene trees with each evolutionary hypothesis.**

| Nanohaloarchaea + Haloarchaea[a] | Nanohaloarchaea + DPANN[b] | Methanantronarchaeia basal to Methanotecta[c] |
|---|---|---|
| 50S_rprotein_3 | 30S_rprotein_1 | 50S_protein_2 |
| 50S_rprotein_5 | endA | dph2 |
| Afung | ftsY | flpA |
| atpA | Hypo_protein_16 | ftsY |
| atpD | Hypo_protein_52 | Hypo_protein_76 |
| atpB | Hypo_protein_76 | iscu2 |
| cysD | Hypo_protein_100 | pcn |
| eif5a | Smli | prf1 |
| glyA | Rnj_1 | Rgy |
| Hypo_protein_113 | rpoK | rnj_1 |
| Hypo_protein_78 | Signal_recog_peptide | rpoA1 |
| Hypo_protein_79 | rpoA | rpoA2 |
| Hypo_protein_83 | rpoB | rpoE1 |
| iscU2 | rps5 | rps7 |
| proS | rps7 | trm1 |
| rps8e | | |

**a:** hypothesis of the Nanohaloarchaea-Haloarchaea sister group, and uses the concordance factor tree in Fig. S11.
**b:** hypothesis of the Nanohaloarchaea falling into DPANN, and uses the concordance factor tree in Fig. S12.
**c:** hypothesis that places the Methanonatronarchaeia at the base of the Methanotecta, and uses the tree in Fig. S12.

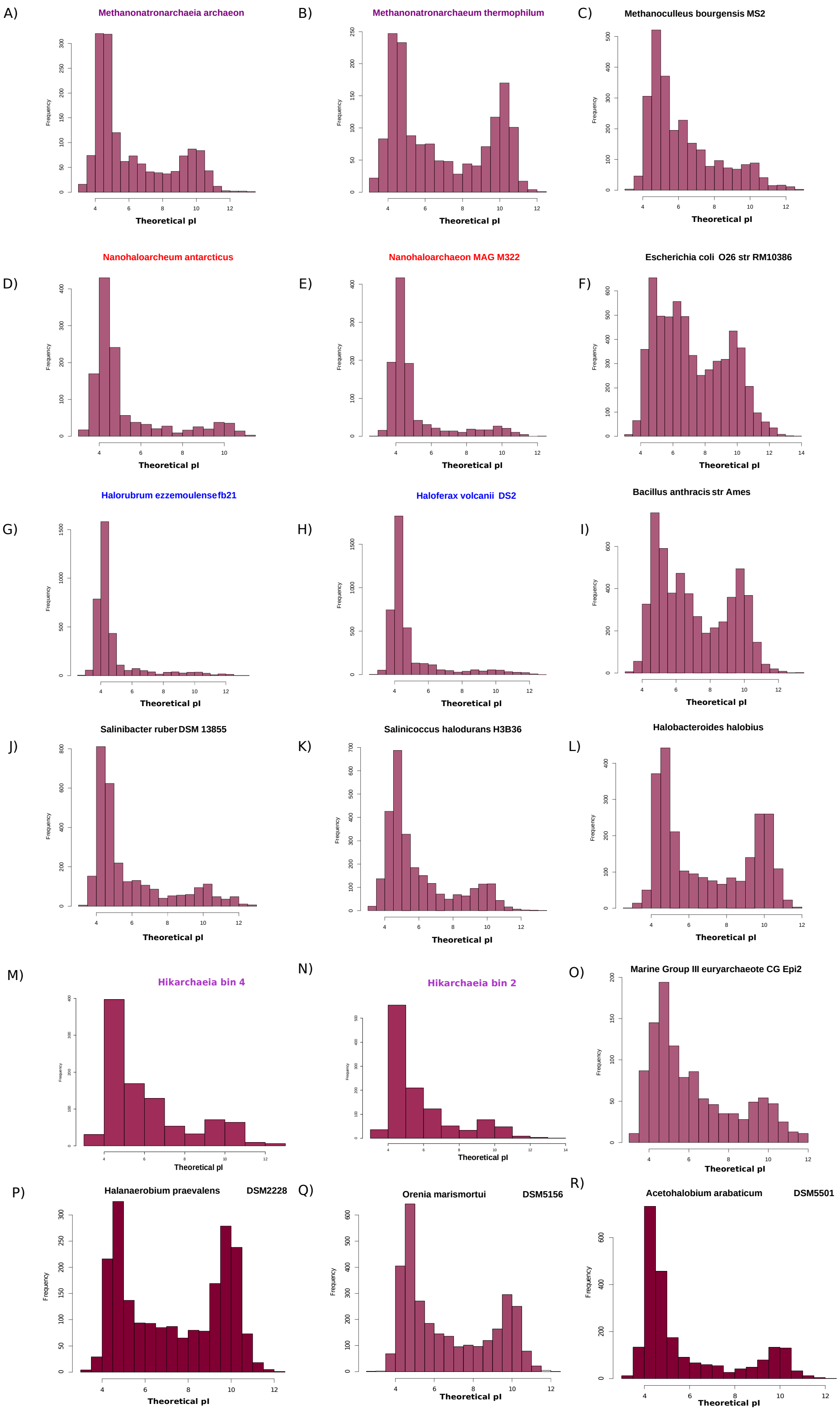
A)
Methanonatronarchaeia archaeon
B)
Methanonatronarchaeum thermophilum
C)
Methanoculleus bourgensis MS2
D)
Nanohaloarcheum antarcticus
E)
Nanohaloarchaeon MAG M322
F)
Escherichia coli O26 str RM10386
G)
Halorubrum ezzemoulensefb21
H)
Haloferax volcanii DS2
I)
Bacillus anthracis str Ames
J)
Salinibacter ruber DSM 13855
K)
Salinicoccus halodurans H3B36
L)
Halobacteroides halobius
M)
Hikarchaeia bin 4
N)
Hikarchaeia bin 2
O)
Marine Group III euryarchaeote CG Epi2
P)
Halanaerobium praevalens DSM2228
Q)
Orenia marismortui DSM5156
R)
Acetohalobium arabaticum DSM5501
Frequency
Theoretical pI

**Fig S13.Protein theoretical isoelectric point distribution in selected proteomes.** Counts (frequency) of proteins at each isoelectric point from selected proteomes. A unimodal distribution, biased towards a lower pI, is indicative of a proteome using the salt-in strategy (Oren, 2008). The Methanonatronarchaeia (A,B) have been reported to use the same strategy (Sorokin *et al.*, 2017), but the intracellular concentrations of potassium have yet to leave a significant impact on pI distribution in their proteomes. These may be similar to the proteomes of the *Halanaerobiales* (P-R), who are salt-in strategists without a bias for acidic amino acids (Bardavid & Oren, 2012). Other users of the salt-in strategy include the Nanohaloarchaea (D,E), Haloarchaea (G,H), *Salinibacter ruber* (J, Bacteroides), and *Salinicoccus halodurans* (K, Firmicutes). For reference are proteomes of those organisms that do not use the salt-in strategy (or where the salt-in strategy has not been experimentally confirmed): C, F, I, L-O. Two Hikarchaeia bins are displayed in M and N, while these organisms are not considered halophiles their proteomes share a similar acidity bias to the Haloarchaea.

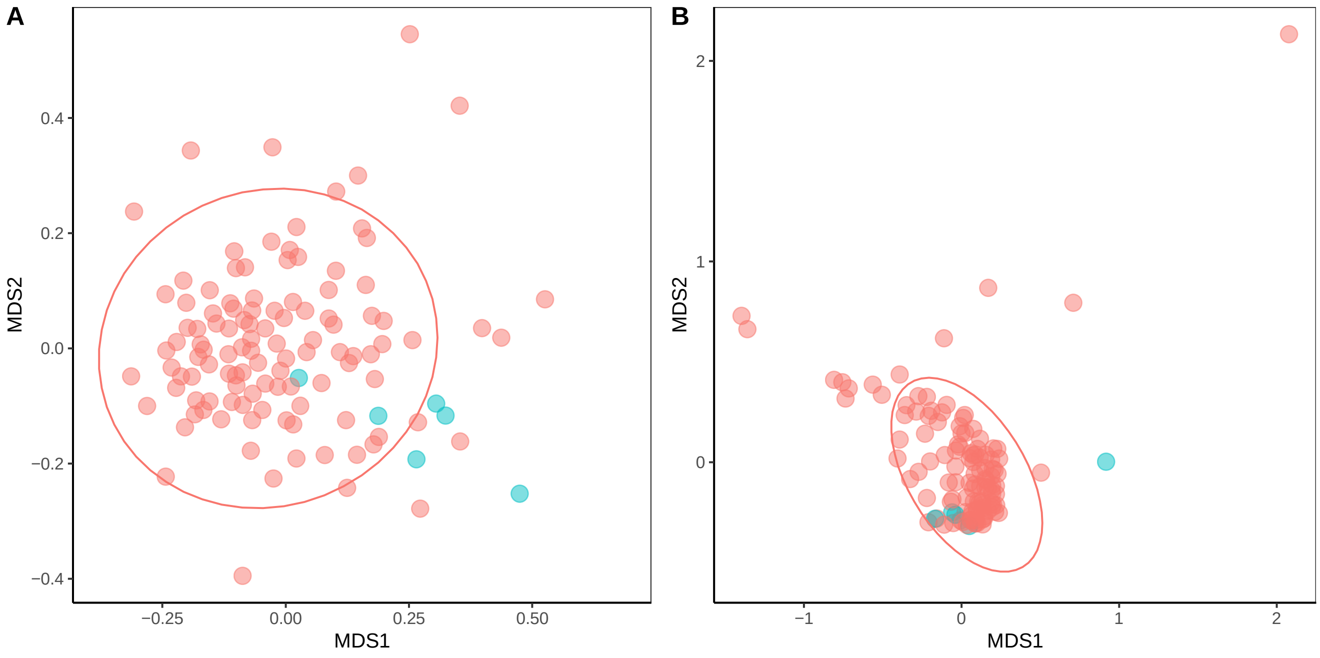


Fig S14. **Clustering of gene families in Thaumarchaeota subsets.** The categorical mantel test was used as a test of significance, where two classes (ATPase gene (in blue) vs. non-ATPase gene) were defined with a 95% confidence ellipse. A) Ordination of gene families in the Thaumarchaeota that have received V-Type ATPases via HGT (Wang et al., 2019). These transfers are detected in this analysis, the categorical mantel test p-value = 0.024. B) Ordination of gene families in neutrophilic Thaumarchaeota that vertically inherited A-Type ATPases. None of the ATPases, apart from AtpI, stands out as atypical, with a p-value = 0.576.
